# TRIM28-dependent SUMOylation protects the adult ovary from activation of the testicular pathway

**DOI:** 10.1101/2021.03.24.436749

**Authors:** Moïra Rossitto, Stephanie Déjardin, Chris M Rands, Stephanie Le Gras, Roberta Migale, Mahmoud-Reza Rafiee, Yasmine Neirijnck, Alain Pruvost, Anvi Laetitia Nguyen, Guillaume Bossis, Florence Cammas, Lionel Le Gallic, Dagmar Wilhelm, Robin Lovell-Badge, Brigitte Boizet-Bonhoure, Serge Nef, Francis Poulat

## Abstract

Gonadal sexual fate in mammals is determined during embryonic development and must be actively maintained in adulthood. In the mouse ovary, oestrogen receptors and FOXL2 protect ovarian granulosa cells from transdifferentiation into Sertoli cells, their testicular counterpart. However, the mechanism underlying their protective effect is unknown. Here, we show that TRIM28 is required to prevent female-to-male sex reversal of the mouse ovary after birth. We found that upon loss of *Trim28*, ovarian granulosa cells transdifferentiate to Sertoli cells through an intermediate cell type, different from gonadal embryonic progenitors. TRIM28 is recruited on chromatin in the proximity of FOXL2 to maintain the ovarian pathway and to repress testicular-specific genes. The role of TRIM28 in ovarian maintenance depends on its E3-SUMO ligase activity that regulates the sex-specific SUMOylation profile of ovarian-specific genes. Our study identifies TRIM28 as a key factor in protecting the adult ovary from the testicular pathway.

---

For long time, it was thought that in mammals, adult gonadal sex assignment was determined and fixed during embryonic development. Any perturbation during this period leads to various disorders of sexual development. However, some teleost fish species display sequential hermaphroditism: gonadal sex is not definitively established in adulthood, and social stimuli can re-assign gonads to the opposite sex (for review see^1^). Moreover, postnatal sex-reversal has been observed in several mouse models: ovarian masculinization upon deletion of oestrogen receptor 1 and 2 (*Esr1-2*)^2^ or of *Cyp19a1*^3^, as well as after postnatal conditional knock-out (cKO) of *FoxL2*^4^ and ectopic ovarian expression of *Dmrt1*^5^. In these cases, the initial cellular event is ovarian-to-testicular transdifferentiation of the supporting cell lineage (granulosa cells to Sertoli cells). Conversely, deletion of *Dmrt1* in postnatal testes^6^ or of both *Sox8* and *Sox9*^7^ induces Sertoli-to-granulosa cell transdifferentiation. These results indicate that granulosa and Sertoli cells retain the ability to transdifferentiate into the opposite sexual fate, and that constant repression of the alternative fate in adult life is required to maintain their cell fate identity and function. However, there is only limited information on the epigenetic and transcriptional programmes implicated in cell fate reprogramming of the supporting lineage.

We previously showed that the epigenetic regulator TRIM28 is a partner of SOX9 in mouse foetal Sertoli cells^8^. TRIM28 is a versatile nuclear scaffold protein that coordinates the assembly of protein complexes containing different chromatin remodelling factors. It can be recruited on chromatin upon interaction with DNA-binding proteins, such as KRAB-ZNF family members^9, 10, 11^, or with transcription factors ^12, 13, 14^. TRIM28 was originally associated with transcriptional repression^9^ and heterochromatin formation^15, 16^; however, many evidences show that it also positively (regulates gene expression^12, 13, 14, 17^ and controls transcriptional pausing^18, 19^. Despite its interaction with SOX9, cKO of *Trim28* in Sertoli cells results in adult males with hypoplastic testes and spermatogenesis defects, but no sex reversal^20^. This suggest that in Sertoli cells, TRIM28 is required to control spermatogenesis, but not for the maintenance of the somatic cell component of the testis.

To understand its role in ovarian physiology, we generated a cKO of *Trim28* in the somatic compartment of the developing mouse ovary. We observed sex reversal in adult ovaries where the follicular structure progressively reorganized in pseudo-tubules with Sertoli-like cells. We then combined mouse genetic with transcriptomic and genomic approaches to determine the molecular action of TRIM28 and its interplay with FOXL2 in adult ovaries. Our data show that TRIM28 maintain the adult ovarian phenotypes through its SUMO-E3 ligase activity that controls the granulosa cells programme and represses the Sertoli cell pathway.

## Results

### Deletion of *Trim28* induces masculinization of adult ovary

Double immunostaining of XX gonads at 13.5 days post-coitum (dpc) showed that TRIM28 is co-expressed with FOXL2 in ovarian pre-granulosa cells, (Fig. S1). To study its role in this crucial ovarian lineage, we generated a mouse line in which *Trim28* can be conditionally deleted using the *Nr5a1:Cre*^21, 22^ transgenic line (*Trim28*^*flox/flox*^; *Nr5a1:Cre* referred as *Trim28*^*cKO*^ or cKO in the text/figures). In 13.5 dpc cKO ovaries, nuclear TRIM28 signal was strongly decreased in FOXL2-positive pre-granulosa cells, whereas it was still present at heterochromatin foci, and was nearly disappeared at E18.8 (fig. S1). At birth, XX cKO mice displayed normal external female genitalia, without any obvious ovarian structure abnormality at 3 days post-partum (dpp) (fig. S2). In FOXL2-positive immature granulosa cells, we did not detect any signal for TRIM28 and SOX8/SOX9, two Sertoli cell markers (fig. S2). Unlike granulosa cells that looked normal at this stage, oocytes were larger, suggesting an early and indirect effect of TRIM28 absence on oogenesis. This suggests that TRIM28 is not required for foetal ovary differentiation. However, as TRIM28 is still expressed in pre-granulosa cells at 13.5 dpc, a potential role in the primary ovarian determination that occurs at ∼11.5 dpc cannot be excluded.

In several follicles of 20 dpp *Trim28*^*cKO*^ ovaries, SOX8 was expressed in groups of cells that stopped expressing FOXL2 (Fig. 1a). Double immunostaining showed that some SOX8-positive cells also expressed SOX9, suggesting that SOX8 expression precedes SOX9, unlike what observed in mouse embryonic testes^23^. As SOX8 and SOX9 are Sertoli cell markers, this suggests that foetal deletion of *Trim28* in pre-granulosa cells might induce their reprogramming towards Sertoli cells after birth, as described for *Foxl2* deletion^4^ and oestrogen receptor double knock-out^2^.

**Fig. 1:**
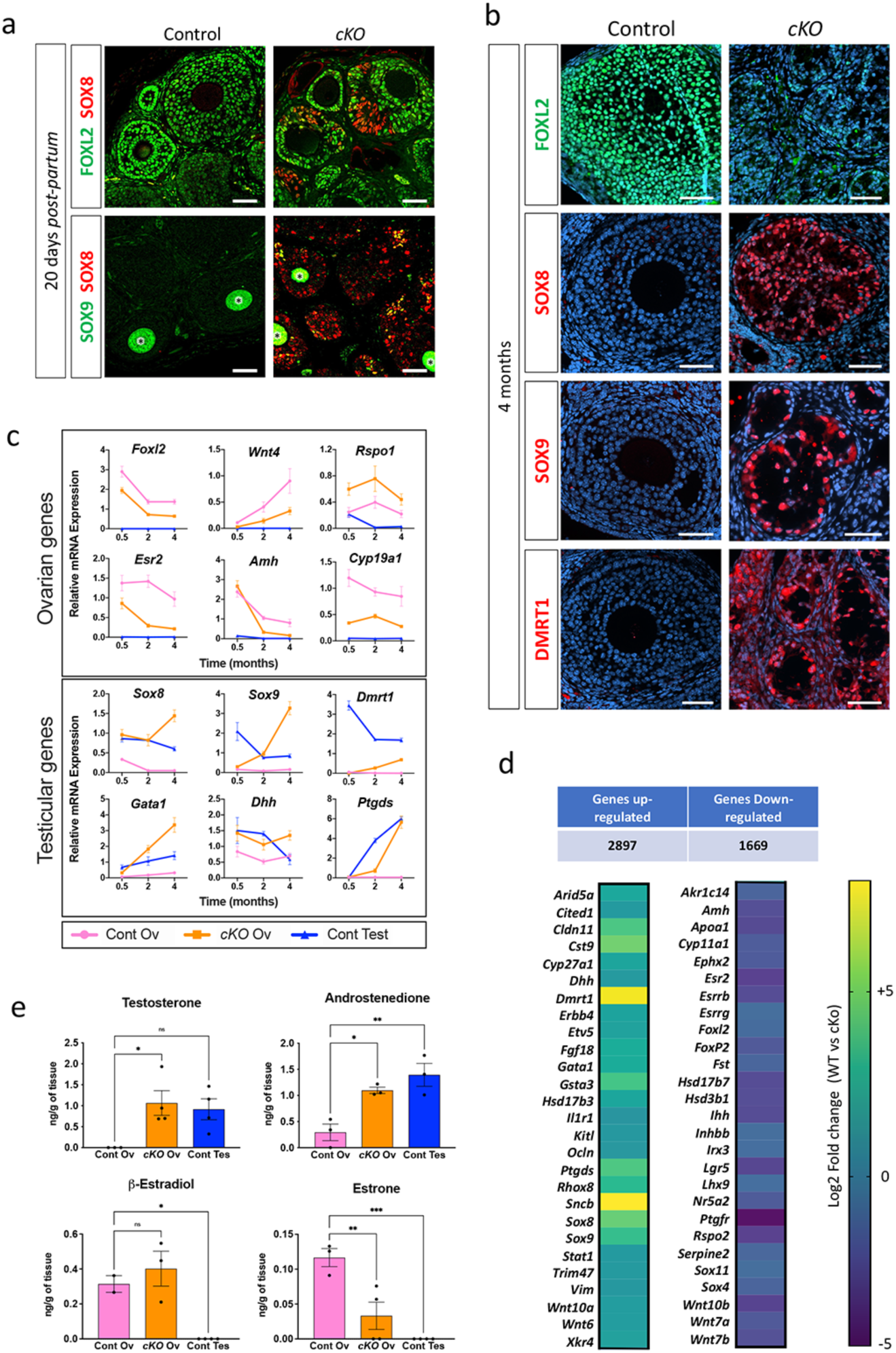
*Trim28* loss in granulosa cells induces masculinization of the adult ovary. **a**, compared with control ovaries, in granulosa cells of 20 dpp *Trim28*^*cKO*^ ovaries, FOXL2 expression is progressively lost and SOX8 (Sertoli cell marker) starts to be expressed. Among the SOX8-positive cells, few express also SOX9, suggesting that SOX8 may precede SOX9. Green staining of oocytes (*) is a non-specific antibody artifact of early folliculogenesis^24^. Scale bar: 50 μm. **b**, in 4-month-old *Trim28*^*cKO*^ ovaries, transdifferentiation to Sertoli cells is complete. Compared with control ovaries, in *Trim28*^*cKO*^ ovaries FOXL2 signal has almost disappeared, and follicles are reorganized in pseudo-tubules that express the Sertoli markers SOX8, SOX9, and DMRT1. Protein (green or red) are merged with DNA stain (blue). Scale bar: 50 μm. **c**, RT-qPCR analysis of the temporal (in months) gene expression variations in control ovaries (Cont Ov), *Trim28*^*cKO*^ ovaries (*cKO* Ov), and control testes (Cont Test). In *Trim28*^*cKO*^ ovaries, typical ovarian genes are progressively downregulated, but for *Rspo1*, and testis genes are upregulated. Details of the statistical analysis are provided in Source data file. **d**, Heatmap of the RNA-seq analysis of 7-month-old ovaries (see Data S1) showing that 2,896 and 1,669 genes are up- and down-regulated, respectively, in *Trim28*^*cKO*^ compared with control ovaries. Normalized expression values are expressed as Log2 fold change (Control *vs cKO*), from -5 (deep violet) to +8 (yellow). **e**, *Trim28* cKO induces the masculinization of the ovarian steroid profile. Steroids were extracted from 7-month-old control (Cont Ov) and *Trim28*^*cKO*^ (cKO Ov) ovaries, and control testes (Cont Test) and quantified (ng/g of tissue) by mass spectroscopy. Data are the mean ±SEM (n=3 or 4, detailed in Source data file); *\*\*\* P*<0.005, *\*\*P*<0.05, *\* P*<0.05 (One-way ANOVA with Tukey’s multiple comparisons test).

In 8-week-old *Trim28*^*cKO*^ mice, ovarian organization was profoundly changed. Medullar follicles had almost completely lost FOXL2 expression, expressed SOX8 and SOX9, and were reorganized into pseudo-tubular structures, indicative of a process of testis cord formation (fig. S3a, d and g). We never detected any cell that expressed both SOX8 (fig S3b) or SOX9 (fig S3e) and FOXL2, but many cells that expressed both SOX proteins (Fig S3h). Their distribution suggested (like in 20 dpp *Trim28*^*cKO*^ ovaries) that SOX8 might precede SOX9. Conversely, the cortical region presented a less advanced phenotype: as observed in 20 dpp *Trim28*^*cKO*^ ovaries, follicles were still organized, but remodelling had started with groups of cells that stopped expressing FOXL2 and expressed SOX8 and/or SOX9 (fig S3c, f and i). These results show that in *Trim28*^*cKO*^ ovaries, the granulosa-to-Sertoli cell transdifferentiation starts in follicles located in the medulla and then spread to the cortical regions.

In parallel, using the Terminal deoxynucleotidyl transferase dUTP nick end labelling (TUNEL) assay, we did not observe any significant increase in apoptosis in 20 dpp and 8-week-old *Trim28*^*cKO*^ ovaries (fig S4), as previously described for the cKO of *Foxl2*^4^. This excluded the replacement by neo-formed Sertoli cells of granulosa cells eliminated by widespread apoptosis.

In 4-month-old *Trim28*^*cKO*^ females, the transdifferentiation of granulosa cells into Sertoli cells was complete: FOXL2 expression has disappeared, and follicles were completely remodelled into tubular structures with cells that expressed the Sertoli cell markers SOX8, SOX9 and DMRT1 (Fig. 1b). Histological analysis confirmed the progressive reorganization of ovarian follicles into tubular structures and the transdifferentiation of granulosa cells into cells with a Sertoli cell morphology (fig. S5). This reorganization was undetectable in 4-week-old *Trim28*^*cKO*^ ovaries but was clearly visible in the medulla at 8 weeks and was completed at 17 weeks. Germ cells (oocytes) were relatively normal in ovaries with a preserved follicular structure but started to degenerate during transdifferentiation. In 8-week-old ovaries in which the medullar part was reorganized into pseudo-tubules, oocytes had disappeared or were degenerating (fig. S5), and in 17-week-old ovaries they had disappeared.

A recent study showed that *Trim28* hemizygosity affects spermatogonial stem cells and induces testis degeneration^25^. However, we did not observe any change in FOXL2 immunostaining in ovaries from wild type and heterozygous *Trim28*^*cKO*^ mice at the different stages we analysed (fig S6a). Similarly, we did not detect any expression change of the three Sertoli markers *Sox8, Sox9* and *Dmrt1* in heterozygous 3-month-old ovaries (fig S6b). Therefore, the loss of a single *Trim28* allele does not cause transdifferentiation of granulosa cells.

We next examined the temporal expression of several genes with roles in testicular and ovarian sex determination in 0.5- (15 dpp), 2 and 4-month-old ovaries. Reverse transcription quantitative real-time polymerase chain reaction (RT-qPCR) analysis revealed that in *Trim28*^*cKO*^ ovaries, the mRNA level of most ovarian-specific genes was decreased, with the exception of *Rspo1* (Fig. 1c, panel Ovarian genes). Conversely, testicular-specific genes were progressively upregulated (Fig. 1c, panel Testicular genes), confirming the histology and immunofluorescence observations. The expression level of some ovarian (*Foxl2, Esr2, Cyp19a1*, and *Rspo1*) and testicular genes (*Sox8* and *Dhh*) was already modified soon after birth (15 dpp), before changes in *Sox9* and *Dmrt1* and before the detection of histological defects (fig S5).

Bulk RNA-seq experiments using 7-month-old *Trim28*^*cKO*^ ovaries (Data S1), in which transdifferentiation was completed, showed that 1669 genes were significantly downregulated in the absence of *Trim28*, among which 71% are normally expressed in adult granulosa cells^26^, including genes involved in ovarian determination (Fig. 1d, right). Repression of the granulosa cell transcriptome was accompanied by upregulation of 2897 genes that included typical Sertoli and Leydig cell markers (Fig. 1d, left), showing that *Trim28* cKO induces the ovarian transcriptome masculinization. We concluded that *Trim28* deletion in foetal pre-granulosa cells induces the postnatal remodelling of the ovarian transcriptome, leading to its masculinization. Moreover, we observed an important deposition of extracellular matrix around pseudo-tubules (fig. S5) and the upregulation of several genes that encode component of the testicular basal lamina: *Col4a3, Col9a3, Col13a1, Col28a1*, and *Lamc2* (encoding laminin gamma 2).

As several genes involved in steroidogenesis displayed a modified profile (Fig. 1c, fig. S7a-b for temporal analysis, and RNA-seq data, respectively), we used mass spectroscopy to quantify the production of major steroid hormones in control and *Trim28*^*cKO*^ ovaries and control testes from 7-month-old animals (Fig. 1e). Androgen levels (testosterone and androstenedione) in *Trim28*^*cKO*^ ovaries and control testes were similar. Among the oestrogens produced in *Trim28*^*cKO*^ ovaries, estrone was strongly reduced, whereas 17ß-estradiol levels were comparable to those in control ovaries. This can be explained by the persistent expression of *Cyp19a1* (the gene encoding the aromatase that catalyses 17ß-estradiol production) in *Trim28*^*cKO*^ ovaries (Fig. 1c) and by the modified expression of genes encoding hydroxysteroid dehydrogenases (HSD) (Fig. S7a-b). Overall, our results indicate that foetal *Trim28* deletion induces the masculinization of the steroid production profile in adult ovaries.

### Granulosa-to-Sertoli transdifferentiation occurs through an unknown cellular intermediate

To better describe the transdifferentiation process, we performed single-cell RNA sequencing (scRNA-seq) to compare the transcriptomic atlas of gonadal cell types in *Trim28*^*cKO*^ ovaries, control ovaries, and control testes. We analysed 8-week-old gonads because our data (fig. S3) indicated that at this stage, *Trim28*^*cKO*^ ovaries contain a mixed population of Sertoli-like cells and apparently normal granulosa cells. Using the 10X Genomics Single Cell Gene Expression system, we analysed 7,292 cells from *Trim28*^*cKO*^ ovaries, 7,051 from control ovaries, and 42,720 from control testes (total=57,063 cells). A larger number of testis cells was required to sample an equivalent number of testicular somatic cells alongside the abundant spermatogenic cells. We catalogued the different cell populations present in all samples (fig. S8a) based on the expression of known markers (fig. S8b). We confirmed the substantial decrease of *Trim28* expression in *Trim28*^*cKO*^ ovarian cells (fig. S9). In control gonads, we detected the expected cell types, including supporting (granulosa/Sertoli), steroidogenic (theca/Leydig), stroma, spermatogenic, endothelial, immune and blood cells (fig. S8), consistent with previous single-cell transcriptomic studies of adult mouse/human testis/ovaries^27, 28, 29^. We then focused on the supporting cell lineages. We identified 3,106 supporting cells that expressed granulosa and/or Sertoli cell markers (n=1,112 in *Trim28*^*cKO*^ ovaries, n=1,446 in control ovaries, and n=548 in control testes) (Fig. 2a). In *Trim28*^*cKO*^ ovaries, transcriptional profiles were asynchronous, some supporting cells were grouped with control granulosa cells and expressed *Esr2, Amh, Foxl2, Wnt4, Hsd17b1*, and *Nr5a2*, indicating that they still had a granulosa-like transcriptome (Fig. 2b). However, we also observed a gradient of gene expression from granulosa to Sertoli cells *via* some intermediate *Trim28*^*cKO*^ ovarian supporting cells (Fig. 2a) that expressed some Sertoli markers at various levels and at different stages of transdifferentiation.

**Fig. 2:**
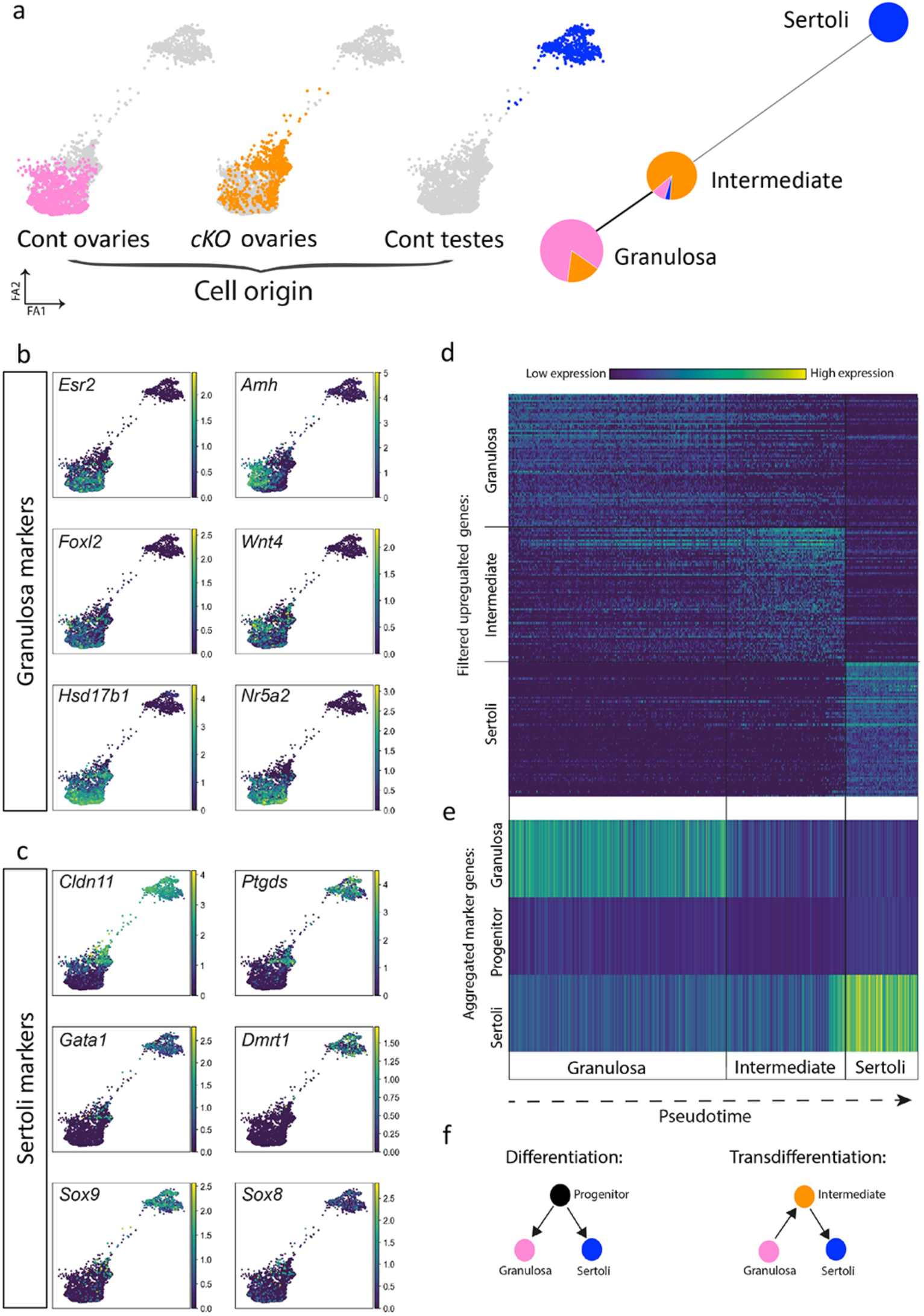
scRNA-seq analysis of ovarian and testis supporting cells reveals an intermediate cell population during transdifferentiation. **a**, Force directed graphs showing the scRNA-seq results of adult *Trim28*^*cKO*^ ovarian supporting cells (orange), control granulosa (pink), and Sertoli cells (blue) (left). Each dot is one cell (coloured according to the sample of origin), and the distance between cells indicates their inferred transcriptional similarity. Leiden clustering divided the cells in three populations displayed using partition-based graph abstraction (right). Each node represents a cell cluster, and the proportion of *Trim28*^*cKO*^ and control granulosa and Sertoli cells is shown as a pie chart on each node. The edges between nodes represent the neighbourhood relation among clusters with a thicker line showing a stronger connection between clusters. **b and c**, Gene expression of selected granulosa and Sertoli cell markers in the supporting cells analysed in (**a)**. Each dot corresponds to one cell from (**a)**, and gene expression level ranges from 0 (purple) to high (yellow). **d**, Heatmap showing the expression level of the top filtered differentially expressed genes in the three cell clusters along the pseudo-time. See Table S3 for the full list of genes. **e**, Heatmap showing the mean expression levels in the three cell clusters along the pseudo-time of several thousand genes from a previous study on the granulosa, supporting progenitor, and Sertoli cell lineages^30^. **f**, Schematic illustrating the processes of differentiation and transdifferentiation.

For example, *Cldn11* and *Ptgds* were expressed earlier during transdifferentiation and in more cells, compared with *Gata1, Dmrt1, Sox9* and *Sox8* (Fig. 2c).

We then asked whether these intermediate cells resembled embryonic XX or XY supporting cell progenitors^30^ that de-differentiated from granulosa cells before differentiating into the Sertoli lineage. We aligned all single cells along a pseudo-time (Fig. 2d, 2e, fig. S10)^31^, and divided them in three clusters based on their transcriptional profiles (Fig. 2a, right). This allowed us to identify genes that were upregulated in the granulosa, intermediate, and Sertoli cell populations (Fig. 2d, Data S3). Analysis of the mean expression of 1,743 supporting progenitor cell markers^30^ showed that they were weakly expressed in intermediate cells (Fig. 2e). This indicated that this population was distinct from embryonic progenitors. Gene Ontology enrichment analysis of the genes expressed in the intermediate population gave only general terms, such as “response to stimulus”, “cell death”, and “cell differentiation” (Data S4). Overall, the scRNA-seq analysis showed that in adult ovaries, *Trim28* cKO leads to transdifferentiation of the supporting lineage from the granulosa to the Sertoli cell fate. Moreover, granulosa cells do not transdifferentiate into Sertoli cell by returning to an embryonic progenitor state, but via a different and novel cell intermediate (Fig. 2f).

### TRIM28 acts in concert with FOXL2 on chromatin

As the *Trim28*^*cKO*^ phenotype was similar to that of mice after *Foxl2* deletion in adult ovarian follicles^4^, we asked whether these two proteins co-regulated common target genes in the ovary. Immunofluorescence analysis confirmed that TRIM28 and FOXL2 were strongly co-expressed in the nucleus of adult control follicular granulosa cells and to a lesser extent in theca stromal cells. Both were almost undetectable in *Trim28*^*cKO*^ ovaries (Fig. 3a). Next, we performed TRIM28 and FOXL2 chromatin immunoprecipitation (ChIP) followed by next-generation sequencing (ChIP-seq) in control ovaries to gain a global view of TRIM28 and FOXL2 co-localization genome-wide. Comparison of the heatmaps of their co-binding to chromatin (Fig. 3b) showed that in ovaries, FOXL2 ChIP-seq reads strongly mapped to regions occupied by TRIM28 (Fig. 3b, blue panel). Similarly, TRIM28 ChIP-seq reads strongly mapped to FOXL2 peaks (Fig. 3b, red panel). Analysis of the overlap between TRIM28 and FOXL2 peaks confirmed that these proteins shared common genomic targets (62 and 55% respectively, Fig. 3b Venn diagram). TRIM28 and FOXL2 bound to overlapping regions of genes that have a central role in ovarian determination, such as *FoxL2, Esr2, Fst* (Fig. 3c), and of genes expressed in granulosa cells (Fig. S11). As these genes were downregulated in *Trim28*^*cKO*^ ovaries, this suggests that TRIM28 and FOXL2 positively regulate major granulosa cell genes. For instance, *Wnt4*, which was downregulated in *Trim28*^*cKO*^ ovaries (Fig 1c), displayed several TRIM28 and FOXL2 peaks in control ovaries (Fig. S11). Conversely, *Rspo1*, which is upstream of *Wnt4* in the ovarian-determining cascade^32^, was upregulated in *Trim28*^*cKO*^ ovaries (Fig 1c). Analysis of the TRIM28/FOXL2 genomic profiles did not highlight any binding on *Rspo1* (fig. S11), suggesting that its regulation in the adult ovary is independent of TRIM28 and FOXL2. Moreover, in the absence of TRIM28, *Wnt4* expression seems to be independent from *Rspo1* expression level.

**Fig. 3:**
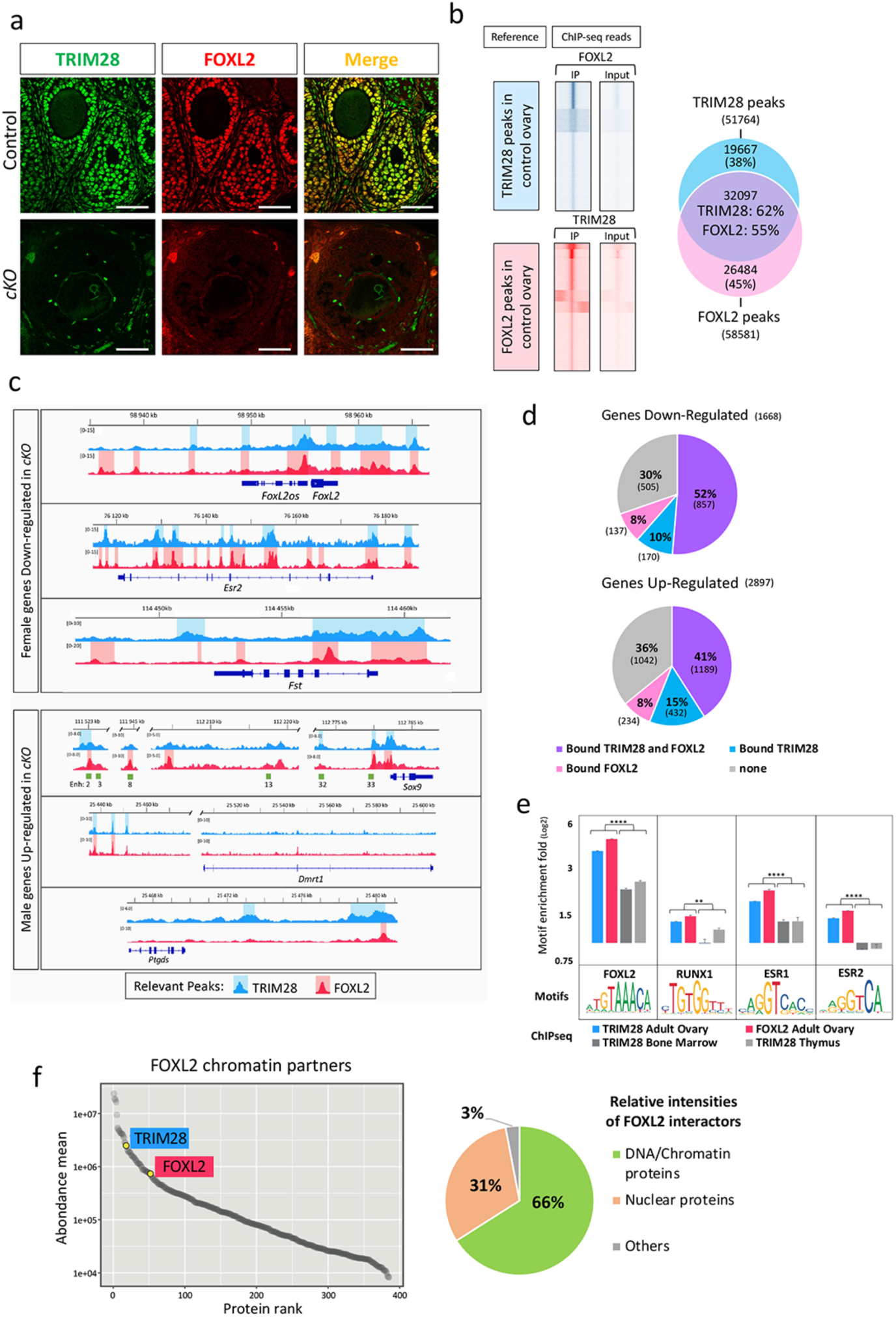
TRIM28 and FOXL2 act together on chromatin to maintain the ovarian pathway. **a**, TRIM28 and FOXL2 are co-expressed in the nucleus of most follicular granulosa cells in 4-month-old control ovaries and in cells with flat nucleus surrounding follicles (identified as steroidogenic theca cells). In *Trim28*^*cKO*^ ovaries, only few cells expressed FOXL2. Scale bar: 10 μm. **b**, Overlap between TRIM28 and FOXL2 genomic localization in the adult ovary. Heatmaps in blue represent FOXL2 ChIP-seq and inputs reads mapped on TRIM28 peaks (± 1kb from the centre). Red traces represent TRIM28 ChIP-seq and inputs reads mapped on FOXL2 peaks. The Venn diagram on the right shows that 32,097 of the 51,764 TRIM28 peaks (62%) and of the 58,581 FOXL2 peaks (55%) overlap in control ovaries. **c**, Examples of TRIM28 and/or FOXL2 peaks in/around genes the expression of which is altered in *Trim28*^*cKO*^ ovaries. Upper panel: ovarian-specific genes downregulated in *Trim28*^*cKO*^ ovaries (see also fig. S11). The *FoxL2* gene is represented with the co-regulated non-coding *FoxL2os* gene ^34^. Lower panel: testicular-specific genes upregulated in *Trim28*^*cKO*^ ovaries (see also fig. S12). Green rectangles in the *Sox9* panel: putative enhancers that with enhancer 13, are crucial for sex determination^33^. Relevant ChIP-seq peaks are highlighted in light blue (TRIM28) and light red (FOXL2). **d**, Pie charts showing up- and down-regulated genes in *Trim28*^*cKO*^ ovaries that are bound by TRIM28 and/or FOXL2. Genes are listed in Data S7. **e**, Enrichment for binding motifs of transcription factors involved in granulosa cell fate maintenance (FOXL2, RUNX1 and ESR1/2) in reads of TRIM28 and FOXL2 ChIP-seq of adult control ovaries (this study), and TRIM28 ChIP-seq of bone marrow^35^ and of thymus^36^. **f**, Left, plot showing enriched proteins, ranked by relative abundance, identified by FOXL2 ChIP-SICAP. Only significant proteins (>2-fold enrichment over No antibody control, n=2) are shown. TRIM28 was identified amongst the top 20 proteins found to interact with FOXL2. Pie-chart (right) shows the percentage of the relative intensities of FOXL2 chromatin partners, normalized to the total abundance of the enriched proteins.

Of note, 52% of the genes downregulated in *Trim28*^*cKO*^ ovaries interacted with TRIM28 and FOXL2 in control ovaries (Fig. 3d). Similarly, many testicular-specific genes upregulated in *Trim28*^*cKO*^ ovaries were bound by TRIM28 and FOXL2 (41%, 1,189 of 2,897), suggesting that TRIM28 and FOXL2 may have a repressive effect on the transcriptional activities of these genes in wild type ovary (Fig. 3d and fig. S12). For example, within the 2-Mb gene desert surrounding the *Sox9* gene, TRIM28 and/or FOXL2 peaks were in close proximity of some of the many enhancers implicated in gonadal *Sox9* expression regulation^33^, and also in the proximal promoter and gene body (Fig. 3c, lower panel). Similarly, the distal upstream regions of *Dmrt1* and *Ptgds*, which are both upregulated in *Trim28*^*cKO*^ ovaries, displayed overlapping regions of TRIM28 and FOXL2 binding (Fig. 3c, lower panels), like other genes, such as *Cldn11* that is expressed in Sertoli cells and upregulated in *Trim28*^*cKO*^ ovaries (fig. S12).

We also analysed DNA motif enrichment for the binding sites of the major granulosa-specific transcription factors (FOXL2^4^, RUNX^22^ and ESR1/2^2^) in TRIM28 and FOXL2 ChIP-seq data, as previously described^8^. We observed a significant enrichment for these motifs in regions bound by TRIM28 and FOXL2 in the ovary compared with regions bound by TRIM28 in bone marrow^35^ and thymus^36^ (Fig. 3e). This shows that in adult ovaries, both TRIM28 and FOXL2 bind to regions that display a genomic signature with binding sites for major ovarian-specific transcription factors.

To confirm that TRIM28 and FOXL2 co-localized on chromatin, we performed FOXL2 ChIP and selective isolation of chromatin-associated proteins (ChIP-SICAP) followed by mass spectrometry that provides only information relative to on-chromatin interactions^37^. We obtained a list of proteins co-localized with FOXL2 on ovarian chromatin that we ranked by their relative abundance. TRIM28 was amongst the top 20 FOXL2 interactors, confirming that it is recruited on chromatin regions very close to FOXL2 (Fig. 3f, left). It should be noted that TRIM28 has been recently shown^38^ to interact with chromatin through two regions of the RBCC domains^39^ (amino acids 298 to 305, and 349 to 366) and an intrinsically disordered region (amino acids 555 to 591). A gene ontology analysis of the protein list (that will be analysed and published elsewhere) showed that these proteins were mainly nuclear and chromatin factors, with only 3% of potential contaminants, demonstrating the technique specificity (Fig. 3f, right). These results are supported by a previous proteomic analysis of murine granulosa and pituitary-derived cell lines showing that TRIM28 and FOXL2 are engaged in common protein complexes^40^. Overall, the previous data on FOXL2^4^ and our results show that in the ovary, TRIM28 and FOXL2 are implicated in the same genetic pathway to maintain the ovarian cell fate. On chromatin, this is achieved through their colocalization on regulatory regions of genes that control the granulosa and Sertoli cell fates. Our data suggest that the TRIM28 /FOXL2 pathway supports the granulosa cell fate by maintaining the ovarian identity and suppressing the testicular identity.

### Mutation of the SUMO-E3 ligase activity of TRIM28 recapitulates the lack of TRIM28 in granulosa cells

TRIM28 acts as a SUMO-E3 ligase by interacting with the SUMO-E2 conjugating enzyme UBC9 (encoded by the *Ube2i* gene) via the Plant homeodomain (PHD) and can self-SUMOylate^41^ (Fig. 4a). SUMOylation is involved in transcriptional regulation and regulates positively or negatively the transcriptional activation capacity and/or stability of many transcription factors, such as FOXL2^42^, ESR2^43^, GATA4^44^, PPARγ and RXR^45^, and of many chromatin-associated proteins^46^. It is also an important histone modification (for review see^47^). Moreover, it has been reported that the SUMOylation status of transcription factors, such as NR5A1^48^, and of androgen receptor^49^ regulates their function in a tissue-specific fashion. Other proteins, such as PCNA^50^, CDK9^51^, NPM1/B23^52^, IRF7^53^, VPS34^54^, α-synuclein, and tau^55^, also are SUMOylated in a TRIM28-dependent manner. To study *in vivo* the role of TRIM28-dependent SUMOylation, we generated a point mutation in exon 13 of mouse *Trim28* within the PHD domain (C651F) that abrogates its SUMO-E3 ligase activity^52^ (fig. S13). *Trim28*^*C651F/+*^ heterozygous mice reproduced normally and did not show any obvious phenotype. However, as we never obtained homozygous mutants when mating heterozygous animals, the homozygous *Trim28*^*C651F*^ mutation (termed *Trim28*^*PHD*^) might be embryonic lethal, like *Trim28* ablation^56^. As heterozygous *Trim28 cKO* (*Nr5a1:Cre*;*Trim28*^*flox/+*^) mice have no phenotype (Fig S6), we generated *Nr5a1:Cre*;*Trim28*^*C651F/flox*^ mice (*Trim28*^*PHD/cKO*^). First, we showed that the TRIM28^C651F^ mutant protein was effectively produced and localized in the nucleus in *Trim28*^*PHD/cKO*^ mutant ovaries (fig. S14). RT-qPCR analysis of 8-week-old ovaries (Fig. 4b) showed that *Trim28* mRNA level in *Trim28*^*PHD/cKO*^ ovaries was intermediate between control (*Trim28*^*+/+*^) and *Trim28*^*cKO*^ ovaries, confirming the presence of TRIM28^C651F^ transcripts. Moreover, ovarian- and testicular-specific genes (*FoxL2, Esr2, Wnt4, Hsd3b1, Ihh*, and *Sox9, Sox8, Dmrt1, Gata1, L-Pgds*, respectively) in *Trim28*^*+/+*^ and *Trim28*^*PHD/+*^ ovaries displayed similar expression levels, showing no dominant effect of the mutated allele. Conversely, in *Trim28*^*PHD/cKO*^ ovaries, ovarian genes were strongly downregulated, and testicular-specific genes were upregulated, like in *Trim28*^*cKO*^ ovaries.

**Fig. 4:**
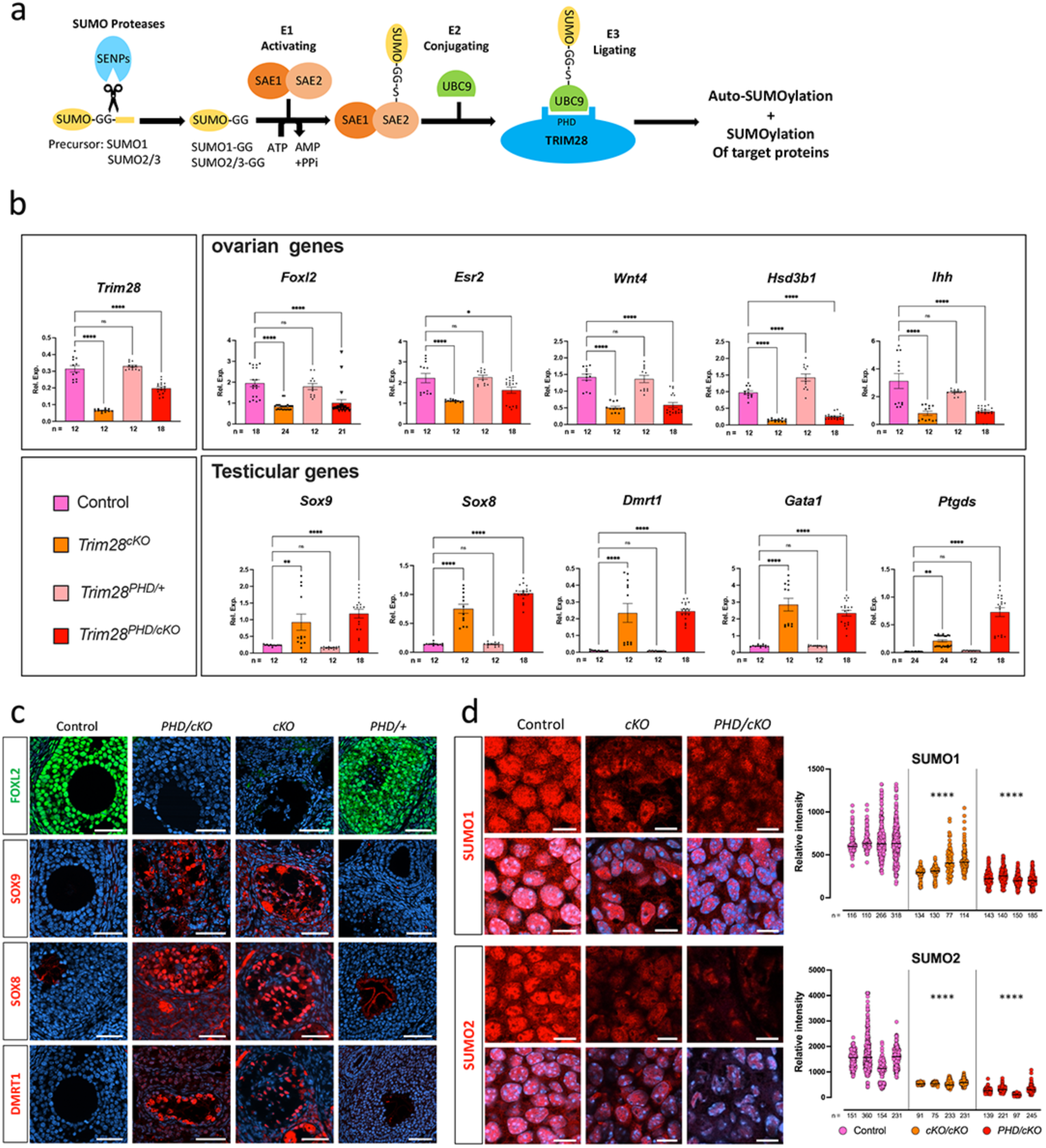
Loss of TRIM28 SUMO-E3 ligase activity in granulosa cells phenocopies *Trim28* conditional knock-out. **a**, Schematic of the SUMO pathway with TRIM28 E3-SUMO ligase activity. After proteolytic maturation by sentrin-specific proteases (SENPs), SUMO C-terminus is activated by the heterodimeric SUMO-activating enzyme E1 (SAE1/SAE2), and then transferred to a cysteine of E2 (UBC9). Subsequently, the E3 ligases (TRIM28) transfer SUMO from E2 to a lysin residue(s) of target proteins. SUMO2 and 3 diverge by only one residue, making them indistinguishable by antibodies, thus they are currently referred to as SUMO2. **B**, RT-qPCR analysis of ovarian- and testicular-specific genes in 8-week-old *Trim28*^*cKO*^, *Trim28*^*PHD/cKO*^, *Trim28*^*PHD/+*^, and control ovaries. Bars are the mean ±SEM, *n* is indicated for each condition. *\*\*\*\*P* <0.0001, *\*\*P*<0.05, (one-way ANOVA with Tukey’s multiple comparisons test). More statistical data are in Source data file. **c**, FOXL2 is expressed in control and *Trim28*^*PHD/+*^ ovaries, but not in *Trim28*^*PHD/cKO*^ and *Trim28*^*cKO*^ ovaries. Like in *Trim28*^*cKO*^ ovaries, SOX9, SOX8 and DMRT1 are expressed in pseudo-tubules of *Trim28*^*Phd/cKO*^ ovaries, but not in control and *Trim28*^*PHD/+*^ ovaries. Protein (green or red) are merged with DNA stains (blue). Scale bar: 50μm. **d**, Confocal microscopy shows strong SUMO1 and 2 nuclear staining in granulosa cells of control ovaries. The staining intensity is markedly decreased in *Trim28*^*cKO*^ and *Trim28*^*PHD/cKO*^ ovaries. SUMO1/2 staining are merged with DNA staining. Right panels: quantification of SUMO1 and SUMO2 signal intensity relative to DNA staining. For the three conditions (control and mutants) each column represent one experiment, *n* represents the number of cells analysed. **** adjusted P value <0.0001 (two-way ANOVA with Dunnett’s multiple comparisons test). More statistical data are in Source data file.

This suggests that *Trim28*^*PHD/cKO*^ and *Trim28*^*cKO*^ ovaries display a similar phenotype. Next, we compared by immunofluorescence analysis, the expression of testis markers (SOX9, SOX8, and DMRT1) and of FOXL2 in *Trim28*^*PHD/cKO*^, *Trim28*^*cKO*^, and control ovaries. Like in *Trim28*^*cKO*^ ovaries, FOXL2 expression was undetectable, whereas we observed expression of the Sertoli cell markers SOX9, SOX8 and DMRT1 within structures organized in pseudo-tubules in *Trim28*^*PHD/cKO*^ ovaries (Fig. 4c). Histological analysis (fig. S15) also showed a similar tissue organization in *Trim28*^*PHD/cKO*^ and *Trim2*^*cKO*^ ovaries. Altogether, these results indicate that the ovarian pathway maintenance in the adult ovary depends on the E3-SUMO ligase activity of TRIM28.

### TRIM28 mutants display a modified SUMOylation landscape

To determine whether the global SUMOylation level in the nucleus of granulosa cells was affected in *Trim28*^*PHD/cKO*^ and *Trim2*^*cKO*^ ovaries, we used a confocal microscopy quantitative analysis with anti-SUMO1 and -SUMO2/3 antibodies (called here SUMO2 because SUMO2 and 3 cannot be differentiated with antibodies). In both *Trim28*^*PHD/cKO*^ and *Trim28*^*cKO*^ ovaries, SUMO1 and particularly SUMO2 nuclear staining were decreased in ovarian somatic cells (Fig. 4d, left), as confirmed by fluorescence quantification (Fig. 4d, right). This shows that the absence of TRIM28 SUMO-E3 ligase activity in ovarian somatic cells decreased the nuclear level of SUMOylation, confirming the link between TRIM28 and this post-transcriptional modification *in vivo*.

As TRIM28 may SUMOylate some transcription factors or chromatin-associated proteins, we determined whether in the two *Trim28* mutant mouse lines, the SUMOylation landscape was modified genome-wide. Quantitative SUMO1 and SUMO2 ChIP-seq analyses in adult *Trim28*^*PHD/cKO*^, *Trim28*^*cKO*^ and control ovaries identified 249,760 chromatin regions that were SUMOylated by SUMO1 or SUMO2 in control ovaries.

As expected, in *Trim28*^*cKO*^ and *Trim28*^*PHD/cKO*^ ovaries, 7.3% and 5.2% of these peaks, respectively, displayed a significantly lower signal (Log2 FC<-1, AdjP value >0.05) and we designated them as hypo-SUMOylated peaks (Fig. 5a, upper panel, and blue spots in fig. S16). The median size of these peaks was <1 kb (0.875 and 0.959 kb for *Trim28*^*cKO*^ and *Trim28*^*PHD/cKO*^, respectively), but bigger than those obtained with TRIM28 or FOXL2 (0.775 and 0.532 kb, respectively). Of note, the number of hypo-SUMOylated peaks was higher in *Trim28*^*cKO*^ than *Trim28*^*PHD/cKO*^ ovaries (SUMO1+SUMO2 peaks: 18,338 *versus* 12,972), suggesting that the C651F mutation may not completely abolish TRIM28 E3-ligase activity, although it induces granulosa-to-Sertoli cell transdifferentiation, as indicated by the similar phenotype of the two mutants.

**Fig 5:**
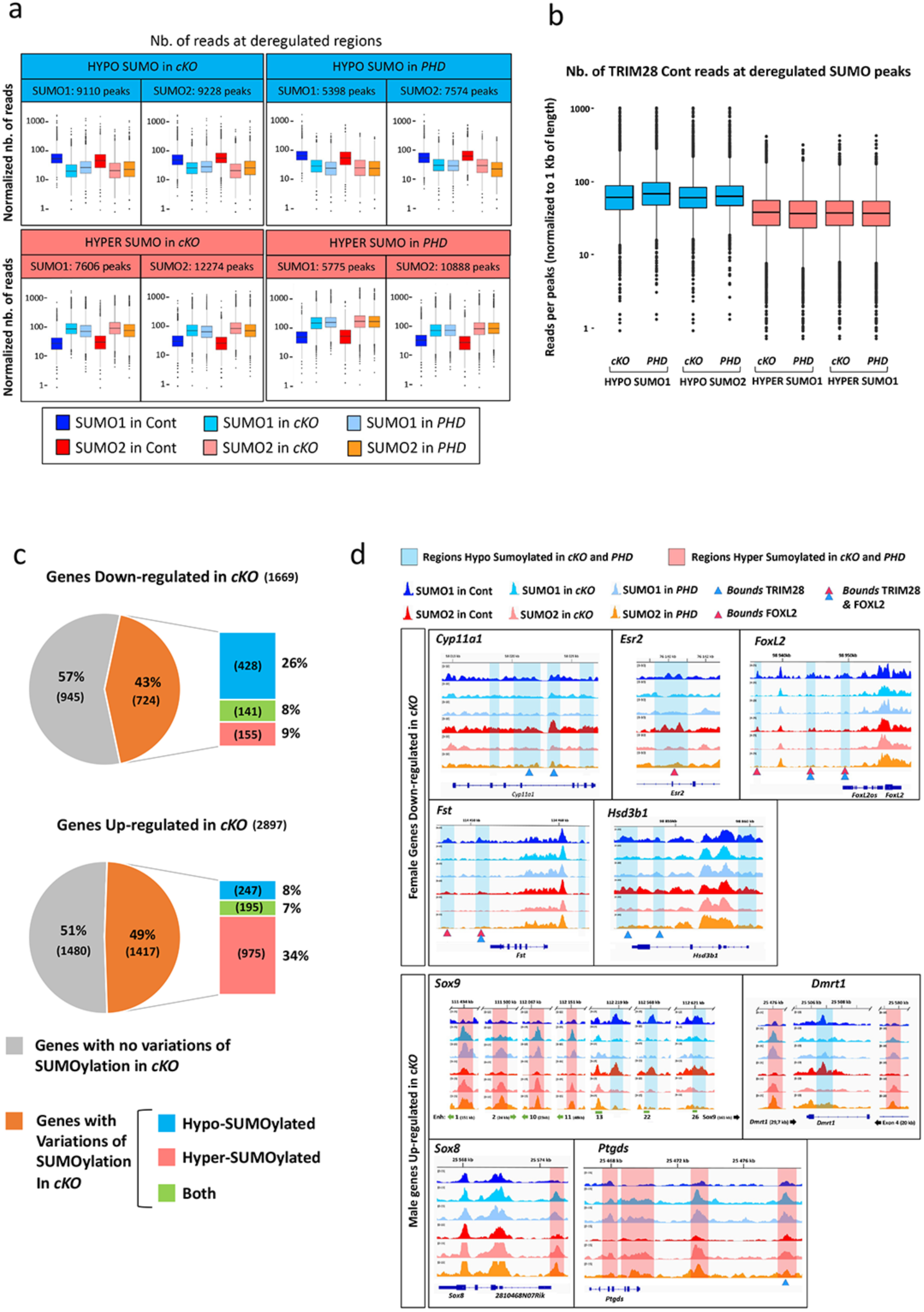
Genome-wide SUMOylation changes in *Trim28*^*cKO*^ and *Trim28*^*Phd/cKO*^ ovaries. **a**, Normalized quantification of SUMO1 and SUMO2 ChIP-seq reads from control (Cont), *Trim28*^*cKO*^ (cKO) and *Trim28*^*Phd/cKO*^ (*PHD*) ovaries mapped on deregulated regions: peaks significantly *decreased* (Log2 Fold Change <1; hypo-SUMOylated; blue), and *increased* (Log2 Fold Change >1; hyper-SUMOylated; red) in *Trim28*^*cKO*^ and *Trim28*^*Phd/cKO*^ ovaries compared with controls. **b**, Normalized quantification of TRIM28 ChIP-seq reads from control (± 1kb from the centre) at SUMO1 and SUMO2 hypo-SUMOylated peaks (blue box plots) and SUMO1 and 2 hyper-SUMOylated peaks (red box plots). cKO: *Trim28*^*cKO*^ *PHD: Trim28*^*Phd/cKO*^. **c**, Pie charts showing that in *Trim28*^*cKO*^ ovaries, downregulated genes with SUMOylation changes are preferentially hypo-SUMOylated, while upregulated genes with SUMOylation changes are preferentially hyper-SUMOylated. Number of genes are between brackets. Genes are listed in Data S7. **d**, Examples of SUMOylation status (SUMO1 and 2) in control, *Trim28*^*cKO*^ (*cKO)*, and *Trim28*^*Phd/cKO*^ (*PHD*) ovaries of genes the expression of which is altered in *Trim28*^*Phd/cKO*^ ovaries. Light blue and red, regions significantly hypo-SUMOylated and hyper-SUMOylated, respectively, in mutants. Blue and red triangles represent the centre of TRIM28 and FOXL2 peaks respectively (see fig 3, S11 and S12).

Quantification of SUMO1 and SUMO2 ChIP-seq reads that mapped to hypo-SUMOylated peaks (Fig 5a) showed that they were markedly decreased in *Trim28*^*cKO*^ and *Trim28*^*PHD/cKO*^ samples (Fig. 5a, upper panel in blue). Moreover, quantification of TRIM28 ChIP-seq reads from control ovaries showed that they mapped strongly to these regions (Fig 5b, box plots in blue). This shows that in control ovaries, TRIM28 occupies chromatin regions that are hypo-SUMOylated in *Trim28*^*cKO*^ ovaries, strongly implying that TRIM28 is the E3-ligase responsible of their SUMOylation in adult ovary (either auto-SUMOylation or SUMOylation of transcription factors located near TRIM28 on chromatin). For example, many hypo-SUMOylated regions in *Trim28*^*cKO*^ ovaries were occupied by TRIM28 and FOXL2 in control ovaries (Fig. S17), suggesting that FOXL2 might be a TRIM28 substrate. Moreover, it has been reported that FOXL2 is SUMOylated in ovarian cell lines^42, 57, 58, 59^ where this modification might promote its stabilization^42, 58, 59^. Similarly, ESR2 stability is regulated by SUMOylation^43^. Analysis of recently published ESR2 ChIP-seq data^60^ also showed that ESR2 peaks overlapped with the hypo-SUMOylated peaks of our mutants, but to a lesser extent than what observed for FOXL2 (fig. S17). RUNX1 is another transcription factor involved in the maintenance of the foetal ovarian fate that shares with FOXL2 a substantial number of genomic targets^22^. Due to the absence of publicly available RUNX1 ChIP-seq data in adult ovaries, we performed SUMOylation assays in cells transfected with wild type TRIM28 or the PHD mutant. We observed that TRIM28 wild type, but not the PHD mutant induced SUMOylation of both FOXL2 and RUNX1 (fig. S18), suggesting that both factors are potential substrates of TRIM28 E3-ligase activity.

However, TRIM28-dependent SUMOylation of transcription factors might also occur before their interaction with chromatin because only a fraction (33 to 45%) of hypo-SUMOylated regions in *Trim28*^*cKO*^ and *Trim28*^*PHD/cKO*^ ovaries were occupied by TRIM28 in control ovaries (fig. S17). We also found a substantial number of SUMO1 or SUMO2 peaks with a significantly stronger signal in *Trim28*^*cKO*^ or *Trim28*^*PHD/cKO*^ than control ovaries (Log2 FC>1, AdjP val >0.05) that we designated as hyper-SUMOylated (Fig. 5b, upper panel, and red spots in fig. S16). ChIP-seq read quantification showed that in *Trim28*^*cKO*^ and *Trim28*^*PHD/cKO*^ ovaries, hyper-SUMOylation (SUMO1 and SUMO2) occurred *de novo* on regions that were less SUMOylated in control ovaries (Fig. 5a, lower red panels). Moreover, quantification of TRIM28 ChIP-seq reads in control ovaries showed that these hyper-SUMOylated regions were poorly occupied by TRIM28 (Fig 5b, box plots in red), unlike hypo-SUMOylated regions (Fig 5b, box plots in blue). In agreement, peak analysis showed nearly no overlap between hypo- and hyper-SUMOylated regions in both mutants (fig. S19). These hyper-SUMOylated peaks might be the signature of Sertoli cell-specific transcription factors expressed in transdifferentiated granulosa cells. To test this hypothesis, we analysed SOX9 and DMRT1 ChIP-seq data during granulosa-to-Sertoli cell transdifferentiation induced by ectopic DMRT1 expression in the ovary^61^. We found that in both *Trim28*^*cKO*^ and *Trim28*^*PHD/cKO*^ ovaries, 14 to 18% of hyper-SUMOylated peaks overlapped with DMRT1 peaks, while 3 to 5% overlapped with those of SOX9 (fig S20). Although more experiments are required to confirm that DMRT1 is SUMOylated, our analysis shows that some hyper-SUMOylated peaks are effectively occupied by DMRT1 and SOX9 during adult reprograming of granulosa to Sertoli cells.

Our results showed that downregulation of the ovarian pathway in *Trim28*^*cKO*^ and *Trim28*^*PHD/cKO*^ ovaries allows the activation of another pathway, inducing the *de novo* SUMOylation of distinct chromatin regions, possibly related to the activated testicular genes. Yet, the RNA-seq analysis of *Trim28*^*cKO*^ ovaries (Data S1) did not highlight the upregulation of any testicular-specific E3-SUMO ligase (e.g. proteins of the PIAS family). This suggests that such ligases are expressed also in granulosa cells.

Analysis of the list of hypo- and hyper-SUMOylated genes highlighted a strong correlation between the very similar phenotypes of the two mutants and gene SUMOylation. Specifically, 5,082 and 4,056 genes were hypo- and hyper-SUMOylated, respectively, in both *Trim28*^*cKO*^ and *Trim28*^*PHD/cKO*^ ovaries (fig. S21a). Some genes showed a mixed SUMOylation pattern (both hypo- and hyper-SUMOylation peaks) (fig. S21b), suggesting a more complex regulation. However, most genes were strictly hypo- (74%) or hyper- (75%) SUMOylated, indicating that they belong to distinct pathways.

Next, we analysed the SUMOylation status of the genes identified as upregulated or downregulated in *Trim28*^*cKO*^ ovaries by RNA-seq. Among the 1,669 downregulated genes (Fig. 5c, upper pie chart), the genes displaying SUMOylation variations were preferentially hypo-SUMOylated (26%), while a minority were hyper-SUMOylated (9%) or both hypo- and hyper-SUMOylated (8%). Ovarian-specific genes that were downregulated in *Trim28*^*cKO*^ ovaries (*Cyp11a1, Esr2, Foxl2, Fst*, and *Hsd3b1*) displayed hypo-SUMOylated peaks in *Trim28*^*cKO*^ and *Trim28*^*PHD/cKO*^ samples for both SUMO1 and SUMO2 (Fig. 5d, upper panels, and fig. S22), where TRIM28 and FOXL2 are bound in control (Fig. 3c).

Conversely, among the testicular genes upregulated in *Trim28*^*cKO*^ ovaries (Fig. 5c, lower pie chart), genes showing SUMOylation variations were preferentially hyper-SUMOylated (34%), and only 8% and 7% were hypo-SUMOylated and both hyper- and hypo-SUMOylated, respectively (examples in Fig. 5d, lower panel, and fig. S23). The key testicular-specific genes *Sox9* and *Dmrt1* that are strongly repressed in granulosa cells showed a mixed SUMOylation pattern in the mutants. At the *Sox9* locus, we observed a mixed hypo- and hyper-SUMOylation pattern in the large regulating region upstream of the gene body: four hyper-SUMOylated peaks and three hypo-SUMOylated peaks in the proximity and along the enhancers 13, 22 and 26^33^. Similarly, in the *Dmrt1* gene, we detected two hyper-SUMOylated regions, one in the gene body and the other upstream, and one hypo-SUMOylated region. These complex SUMOylation patterns could reflect the need of a strict regulation because expression of these two genes must be silenced in granulosa cells. By contrast, *Sox8* and *Ptgds* (like the testicular genes presented in fig. S23) displayed only hyper-SUMOylation peaks, suggesting that SUMOylation might reflect only their transcriptional activation. Another example is *Cldn11*, one of the earliest Sertoli-specific gene (Fig 2c, fig. S10). We detected TRIM28 and FOXL2 peaks at four different regions of the *Cldn11* genomic locus (fig S12), likely to repress its expression. However, the most upstream of these regions, which is an open chromatin region in embryonic gonads^62^, was hyper-SUMOylated in the cKO and PHD mutants (Fig. S23). Therefore, upon disappearance of TRIM28 and/or FOXL2 in mutants, some transcription factors might have access to this potential enhancer, to activate the *Cldn11* gene.

Overall, the TRIM28 E3-ligase controls the maintenance of granulosa cell fate via the specific SUMOylation of ovarian genes. In its absence, a distinct pathway takes place, leading to the hyper-SUMOylation of some Sertoli cell-specific genes that is correlated with their activation.

## Discussion

This study shows that *Trim28* plays a central role in the postnatal maintenance of the ovarian somatic cell fate. Upon *Trim28* loss in foetal pre-granulosa cells, differentiated granulosa cells are reprogrammed, after birth, into Sertoli cells through a previously undescribed intermediate cell type. Therefore, granulosa cells do not dedifferentiate into embryonic progenitors, but acquire a different cell state in which neither ovarian nor testicular master genes are expressed. Moreover, our scRNA-seq and immunofluorescence data confirmed that transdifferentiation is the only possible mechanism of sex reversal and excluded the *de novo* generation of Sertoli cells concomitantly to granulosa cell disappearance (e.g., due to massive apoptosis). Of note, during somatic sex reprogramming in *Foxl2*^*-/-*^ adult ovaries^4^, for ∼1 day following the disappearance of FOXL2 expression, SOX9 cannot be detected, suggesting a similar intermediate step as observed in the present work.

Unexpectedly, structural genes of Sertoli cells, such as *Cldn11*, were upregulated before key genes encoding testicular transcription factors, such as *Sox9* and *Dmrt1*. This suggests that the onset of transdifferentiation might not occur through the activation of a single master gene, such as *Sox9* or *Dmrt1*, but through the global de-repression of the testicular-specific transcriptome. Our observation that TRIM28 is a co-factor of FOXL2 on chromatin supports this hypothesis. In the absence of functional TRIM28, FOXL2 would progressively loose its capacity to repress the testicular pathway, leading to a global de-repression of Sertoli cell genes. The potential role of SOX8 needs to be better investigated. Immunofluorescence, RT-qPCR and scRNA-seq experiments showed that *Sox8* is upregulated before *Sox9* and *Dmrt1* in *Trim28*^*cKO*^ ovaries. However, as SOX8 has a weak trans-activation capacity^23^, the transdifferentiation process might be accelerated by de-repression of testicular-pathway master genes (*Sox9* and *Dmrt1*). A recent study has shown that DMRT1 acts as a pioneering factor required by SOX9 for the optimal activation of its target genes^60^. In our case, the engagement in the testicular pathway might be partial until *Dmrt1* is fully activated. Additional genetic experiments, using double *Trim28* and *Sox8, Sox9* or *Dmrt1* knock-out lines are required to answer this question.

At the organ level, transdifferentiation is first completed in the medulla and then extends to the cortical region. At week 8 post-partum, mutant ovaries displayed medullar pseudo-tubules and cortical follicles: a two-step process also observed in mice where both oestrogen receptors were knocked out^63^. Interestingly, medullar granulosa cells are mostly derived from bi-potential precursors in which primary sex-determination occurs at 11.5 dpc and that are integrated in follicles at puberty^64, 65^. Conversely, cortical follicle pre-granulosa cells are generated mainly by the celomic epithelium from 13.5 dpc until birth and sustain fertility^66, 67^. This suggest that bipotential precursor-derived medullar granulosa cells might be more sensitive to the effect of *Trim28* absence/mutation.

An important finding of our study is the role of TRIM28-dependent SUMOylation in the maintenance of granulosa cell fate. A previous work showed that global SUMOylation of chromatin-associated proteins has a key role in the stabilization of somatic and pluripotent states^68^. Here, we found that TRIM28-dependent SUMOylation, which represents less than 10% of the whole SUMOylation landscape, is sufficient to prevent adult sex-reversal. TRIM28 induces relatively sharp peaks of SUMOylation on chromatin (<1 kb), unlike the large peaks of histone modifications. This might reflect SUMOylated transcription factors. Therefore, a central question is the nature of TRIM28 targets. As TRIM28 can self-SUMOylate^41^, a large number of hypo-SUMOylated peaks in *Trim28*^*cKO*^ and *Trim28*^*PHD/cKO*^ ovary samples may represent TRIM28 SUMOylation; this was confirmed by the overlap between these peaks and TRIM28 peaks in control ovary samples. Similarly, many FOXL2 peaks overlapped with hypo-SUMOylated peaks in *Trim28*^*cKO*^ and *Trim28*^*PHD/cKO*^ ovary samples, and our *in-vitro* data showed that TRIM28 SUMOylates FOXL2. It was previously shown that FOXL2 SUMOylation leads to its stabilization^42 59^. This could explain why in *Trim28*^*cKO*^ ovaries, FOXL2 is undetectable, although the transcript is still present. Indeed, the lack/reduced SUMOylation of FOXL2 might contribute to its progressively decrease/loss in post-natal ovary, leading to transdifferentiation. However, many hypo-SUMOylated peaks did not overlap with TRIM28 or FOXL2, indicating that other transcription factors or chromatin-associated proteins display TRIM28-dependent SUMOylation, particularly FOXL2, ESR1/2 and RUNX1 that are involved in ovarian maintenance ^2, 22^. We also observed that RUNX1 is SUMOylated by TRIM28 *in vitro* and that ESR1 transcriptional activity ^69^ and ESR2 stability^43^ are positively regulated by SUMO-conjugation. Moreover, genome-wide, ESR2 also binds to hypo-SUMOylated peaks, but in a smaller proportion than FOXL2. In the absence of TRIM28, these transcription factors and FOXL2 might lose their capacity to maintain the ovarian programme.

Our data support the hypothesis that a TRIM28-dependent programme of SUMOylation maintains the transcription of ovarian genes and represses genes involved in Sertoli cell fate. These results challenge the dominant view for many years according to which SUMOylation only represses transcription. Our findings and a recent study on the control of adipogenesis by SUMOylation^45^ suggest a more complex scenario: SUMOylation of important cell regulators (e.g. transcription factors) via a specific SUMO-E3 ligase (e.g. TRIM28) might regulate a complete transcriptional programme through activation or repression of target genes.

As the *Trim28*^*cKO*^ and *Trim28*^*PHD/cKO*^ ovarian transcriptomes displayed a strong masculinization, we also observed activation of a *de-novo* pattern of chromatin SUMOylation (i.e. hyper-SUMOylated peaks) that we attributed to the testicular pathway and that is catalysed by a still unknown E3-ligase. These hyper-SUMOylated peaks might represent SUMOylated Sertoli-specific transcription factors, such as DMRT1 or SOX9 that can be SUMOylated^70^. Importantly, by analysing ChIP-seq data obtained by Lindeman and colleagues in ovarian reprograming experiments^60^, we found that a substantial amount of the hyper-SUMOylated peaks from our results co-localized with DMRT1 peaks and to a lesser extend with SOX9 peaks. This shows that both transcription factors are present in hyper-SUMOylated regions and might be SUMOylated (or their partners) independently of TRIM28. SUMO proteomic approaches should answer these questions about hypo- and hyper-SUMOylated peaks.

Altogether, our findings suggest a multi-step model. First, in the absence of TRIM28, FOXL2 that co-localizes on chromatin with TRIM28 would lose its capacity to maintain the expression of granulosa cell-specific genes. Granulosa cells would differentiate into an intermediate state where they express non-sex-specific markers. Second, this would lead to the de-repression of some Sertoli cell-specific genes, such as *Sox8* or *Cldn11*, allowing progressively the induction of strong activators of the Sertoli cell pathway, such as *Dmrt1*. To confirm this model, we need now to generate mice lacking both *Sox8, Sox9* or *Dmrt1* and *Trim28*.

Unlike its role in granulosa cells, TRIM28 is not required for the maintenance of adult Sertoli cells where it is involved in their crosstalk with germ cells^20^ and also in SUMOylation^71^. However, as we could not completely abolish TRIM28 protein expression in pre-granulosa cells before 13.5 dpc, we cannot exclude a role in primary sex-determination that occurs at 11.5 dpc. Indeed, *in vitro* studies have shown that the testis-determining factor SRY, through its interaction with a KRAB-0 protein^72^, might recruits TRIM28 on chromatin to repress ovarian genes^73^. Therefore, more genetic experiments are required to delete *Trim28* using Cre drivers that work earlier, as previously described for *Gata4*^74^.

TRIM28 is an important player in ovarian physiology and therefore, might also have a potential role in genetic diseases causing reproductive disorders. TRIM28 has been recently identified as a tumour suppressor in Wilms’ tumour, a common paediatric kidney malignancy (reviewed in^75^). However, no *TRIM28* mutation has been described so far in patients with reproductive disorders, such as primordial ovarian insufficiency^76^, and in patients with disorders of sexual development (Dr Ken McElrevey personal communication). Besides genetic alterations, environmental factors, such as drugs or chemicals may also interfere with the SUMO-E3 ligase activity of TRIM28 and this could in turn perturb ovarian function and fertility.

