## Supplementary material for "TRIM28-dependent SUMOylation protects the adult ovary from activation of the testicular pathway": methods and supplementary Fig S1 to S23

### Generation of mutant mice

The *Nr5a1:Cre* line [21] and its use to generate conditional knock-outs in ovarian granulosa cells were described previously [22]. The *Trim28* conditional allele was described previously [56]. To generate granulosa *Trim28* conditional knock-out mice, mice carrying *Trim28* loxP-flanked alleles (flox) (*Trim28*$^{flox/flox}$) were crossed with mice bearing the *Nr5a1:Cre* transgene to generate *Trim28*$^{flox/+}$; *Nr5a1:Cre* mice. These mice were then crossed with *Trim28*$^{flox/flox}$ mice to generate *Trim28*$^{flox/flox}$; *Nr5a1:Cre* female mice; these mice were referred to as *Trim28*$^{cKO}$ null mutants. These crosses also generated *Trim28*$^{flox/flox}$ mice without the *Nr5a1:Cre* transgene and *Trim28*$^{flox/+}$; *Nr5a1:Cre* mice; both genotypes did not show any histological defect and were thus referred to as control mice.

The *Trim28*$^{Phd/+}$ mutant mouse line was established at the Mouse Clinical Institute - Institut Clinique de la Souris, Illkirch, France (*http://www-mci.u-strasbg.fr*). *Trim28*$^{Phd/+}$ embryonic stem cells (see fig. S11 for the generation of the targeted C651F mutation in exon 13 of *Trim28*) were used to derive *Trim28*$^{Phd/+}$ mice. To generate granulosa knock-in mice that express only the mutant TRIM28 protein, *Trim28*$^{Phd/+}$ mice and mice bearing the *Nr5a1:Cre* transgene were first crossed to produce *Trim28*$^{Phd/+}$: *Nr5a1:Cre* mice. These mice were then crossed with *Trim28*$^{flox/flox}$ mice to generate *Trim28*$^{Phd/flox}$: *Nr5a1:Cre* mice; these mice were referred to as *Trim28*$^{Phd/cKo}$ knock-in mutants or *PHD* mutants. These crosses also generated *Trim*$^{Phd/flox}$ mice without the Nr*5a1:Cre* transgene, *Trim28*$^{Phd/+}$; Nr*5a1:Cre* mice, and *Trim28*$^{flox/+}$; Nr*5a1:Cre* mice that did not have any histological defect, and were referred to as control mice. No *Trim28*$^{Phd/Phd}$ homozygous mouse was obtained when *Trim28*$^{Phd/+}$ mice were crossed. This suggests that the homozygous PHD mutation may be embryonic lethal, as already observed for the TRIM28$^{HP1box}$ mutation [20]. Animal care and handling were according to the "Reseau des Animaleries de Montpellier" (RAM) guidelines.

*Trim28* floxed allele was genotyped with primers surrounding the loxP insertion site [56]: 5'-GGAATGGTTGTTCATTGGTG-3' and 5'-ACCTTGGCCCATTTATTGATAAAG-3'. The wild type allele gives a PCR product of 152 bp, and the floxed allele of 180 bp.

The *Trim28$^{Phd}$* allele was genotyped with primers that surround the mutation in exon 13 (5'-AAGCCTGTGTTGATGCCTCT-3' and 5' CTTCAGCTACTGGGCCACAC-3') and that give a PCR product of 513 bps in the wild type allele. The mutation introduces a site for the restriction enzyme *Afe*I that cuts the sequences amplified by the primers in two fragments of 240 and 273 bp, respectively. The *Phd* allele gives also a PCR product of 180 bps with the primers used for the *Trim28* floxed allele.

The Nr*5a1:Cre* transgene was genotyped using the 5'-TCGGGGTTTTGTTCTCAGAC-3' and 5'-ATGTTTAGCTGGCCCAAATG-3' primers that give a PCR product of around 500 bp.

PCR conditions for all genotyping were: 94°C for 30sec; 55°C for 30sec, 72°C for 30sec, 30 cycles.

**ChIP-seq analysis.**

Dissected ovaries were snap-frozen and stored at −80◦C. Frozen ovaries were crushed in liquid nitrogen using a mini-liquid nitrogen cooled mortar (Bel-Art ref H37260-0100). Powdered tissue samples were immediately fixed in PBS/1% formaldehyde at room temperature for 20 min. Chromatin shearing was performed using a qSonica Sonicator. Chromatin immunoprecipitation and sequencing libraries were prepared using the Low Cell ChIP-Seq Kit, the Next Gen DNA Library Kit that contains molecular identifiers (MIDs) used to remove PCR duplicates from sequencing data, and the Next Gen Indexing kit (Active motif, ref 104895). For peak calling, the peak caller that best fitted our experimental design and data was chosen and following the ENCODE consortium recommendations[77] (see also: https://www.encodeproject.org/pipelines/).

For TRIM28 and FOXL2 ChIP-seq: Each ChIP-seq library was prepared from a pool of three independent immunoprecipitations (IP), and each IP was prepared with ovaries of two different animals. Reads were mapped to the *Mus musculus* genome (assembly mm10) using Bowtie[78] v1.0.0 with default parameters except for "-p 3 -m 1 –strata –best –chunkmbs 128. BAM files were sorted using SAMtools[79] v0.1.19. The tool rmDupByMids.pl provided by Active Motif was then used to remove duplicated reads. Peaks were called using MACS2[80] with default parameters except for "-g mm -f BAM –broad –broad-cuto 0.1 –keep-dup all".

For SUMO1 and SUMO2 ChIP-seq: Two independent ChIP-seq libraries were prepared. For each library three independent IPs were pooled, each IP prepared from ovaries of two different animals. Reads were mapped to the *Mus musculus* genome (assembly mm10) using Bowtie [78] v1.0.0 with default parameters except for "-p 3 -m 1 –strata –best". BAM files were sorted using SAMtools [79] v0.1.19. The tool rmDupByMids.pl provided by Active Motif was then used to remove duplicated reads. For peak calling data were analysed using the Encode ChIP-seq pipeline v1.3.6[77]. Briefly, the pipeline ran quality controls and called peaks with spp v1.15.5[81]. Spp v1.15.5 was run through the ENCODE pipeline v1.3.6 using the following parameters "-npeak=300000 -fdr=0.01". The fragment length parameter "-speak" was set according to the value estimated by the ENCODE pipeline. Reproductible peaks were kept after the IDR analysis was run. Peaks were annotated relative to genomic features using Homer v4.11.0[82] with Ensembl v92 annotations. Regions differentially bound between conditions were detected as follows. For all proteins of interest, all detected peaks were combined to merge all peaks (249760 peaks) using Bedtools merge v2.26.0. Finally, read counts were normalized across libraries using the method proposed by[83]. Comparisons were performed using the method proposed by [84] implemented in the DESeq2 Bioconductor library (DESeq2 v1.6.3). The resulting p-values were adjusted for multiple testing using the Benjamini and Hochberg method[85]. The following thresholds were used to select differentially bound regions: adjusted p-value $\leq 0.05$, $|\log 2$ Fold-Change$| \geq 1$. Heatmap analyses were performed with SeqMINER v1.3.3g[86]. On all graphs, the presented data are a sub-sample of the initial alignment data (10 million) to make the enrichments in reads comparable. For boxplots, all SUMO peaks were normalized to 1Kb. The number of reads per peak were calculated using bedtools intersect v2.26.0. The graphs are made with R v4.1.1 scripts. Peak overlap was analysed using the intersects tool from the Bedtools package[87] in Galaxy. Motif enrichment was assessed as previously described[8] using the matrix-scan tool (P-value 1e-4) from RSAT[88] (http://rsat.sb-roscoff.fr) and frequency matrices from the JASPAR collection (http://jaspar.genereg.net).

**ChIP-SICAP, mass spectrometry and data analysis**

Ovaries from 8-week-old C57BL/6 mice were dissected in PBS, snap-frozen, and stored at −80 °C. Chromatin was prepared using a modified version of the Active Motif High Sensitivity Chromatin Prep Kit. Frozen tissues were pulverized in liquid nitrogen using the Bel-Art™ SP Scienceware™ Liquid Nitrogen-Cooled Mini Mortar. A pool of ovaries from six mice was used to reach a minimum weight of 50 mg allowing a chromatin yield of at least 60 µg. Samples were fixed in 10 ml of Fixation Buffer (1.5% methanol-free formaldehyde in 1% PBS) on a roller at room temperature for 15 minutes. Fixation was stopped by addition of Stop buffer (Active Motif) for 5 minutes. Chromatin preparation was continued as per the Active Motif protocol except for the sonication steps that were performed in a Bioruptor® Plus sonication device (Diagenode) with the following settings: 40 cycles of 30sec ON/30sec OFF, power "high", constant temperature of 4ºC. ChIP-SICAP experiments were performed in parallel using two biological replicates. ChIP-SICAP allows the identification of chromatin-bound proteins that colocalize with a bait protein (FOXL2 in this study) on DNA. A total of 30 µg of sonicated chromatin was used for each IP using 5 µl of FOXL2 antibody or no-antibody as the negative control. ChIP-SICAP was performed as described by[37]. Briefly after IP, protein complexes were captured on protein G Dynabeads, DNA was biotinylated by Klenow 3'exo in the presence of biotin-7-dATP, followed by TdT in the presence of biotin-11-ddUTP and biotin-11-dCTP, and eluted. Protein-DNA complexes were captured with protease-resistant streptavidin beads [89], formaldehyde cross-linking was reversed, and proteins digested with 300 ng of LysC overnight, followed by 8 hours of incubation with 200 ng trypsin. Digested peptides were cleaned using the stage-tipping technique as follows: digested samples were acidified by addition of 2ul of TFA 10%. For each sample, 50µl of 80% acetonitrile/0.1% formic acid was aliquoted in a LCMS Certified Clear Glass 12 x 32mm Screw Neck Total Recovery Vial (Waters). ZipTip with 0.6 µL C18 resin was pre-treated by pipetting 100% acetonitrile twice by aspirating, discarding and repeating. The ZipTip was equilibrated by pipetting 0.1% TFA, three times by aspirating, discarding and repeating. Acidified samples were then pipetted 10 times with ZipTip to allow peptides to bind to the polymer within the tip. Following tip wash in 0.1% TFA, peptides were eluted in 80% acetonitrile/0.1% formic acid in the glass vial and the eluent dried in a speed vac. Peptides were reconstituted in 8µl of 2% DMSO/0.1% formic acid.

Then, peptides were separated on a 50 cm, 75 µm I.D. Pepmap column over a 70min gradient to be injected into the mass spectrometer (Orbitrap Fusion Lumos) according to the universal Thermo Scientific HCD-IT method. The instrument ran in data-dependent acquisition mode with the most abundant peptides selected for MS/MS by HCD fragmentation and MS/MS by IT. The raw data were analysed using Proteome Discoverer 2.1 (Thermo Scientific). Briefly, Sequest HT node was used to search the spectra using the UniProt *Mus musculus* database. The search parameters were as follows: The precursor mass tolerance was 10 ppm, and MS/MS tolerance was 0.05 Da. Variable modification included Methionine oxidation and N-terminal acetylation. Fixed modification included Cystenine carbamidomethylation. Trypsin and LysC were chosen as the enzymes, and maximum 2 missed cleavages were allowed. Perculator node was used to eliminate false identifications (FDR < 0.01). Protein ratios in Foxl2 assays (n=2) over no-antibody assays (n=2) were calculated in Proteome Discoverer. The quantification values were exported to R studio to analyse significantly enriched proteins using t-test limma package to determine Bayesian moderated t-test p-values and Benjamini-Hochberg (BH) adjusted p-values. We considered proteins with mean fold enrichment > 2 and adj. p-value < 0.1 as enriched proteins.

**RNA isolation, RT-qPCR and RNA-seq analysis**

RNA was extracted from ovaries using TRIZOL (Thermo Fisher Scientific) and processed as described previously[8]. RNA quality was controlled with the Agilent 2100 Bioanalyzer system. RT-qPCR was performed as previously described[8] using the primers listed below and 18S as the reference gene for data normalization. All the statistical analyses were performed using the GraphPad Prism v9 software.

Primers for RT-qPCR (*Gene*: Forward Primer; Reverse Primer), hybridization temperature = 60°C:

*FoxL2*: CGGGGTTCCTCAACAACTC; CATCTGGCAGGAGGCGTA

*Wnt4*: ACTGGACTCCCTCCCTGTCT; TGCCCTTGTCACTGCAAA

*Rspo1*: CGACATGAACAAATGCATCA; CTCCTGACACTTGGTGCAGA

*Esr2*: CCATGATTCTCCTCAACTCCA; TGTCAGCTTCCGGCTACTCT

*Amh*: GGGGAGACTGGAGAACAGC; AGAGCTCGGGCTCCCATA

*Cyp19a1*: GAGAGTTCATGAGAGTCTGGATCA; CATGGAACATGCTTGAGGACT

*Sox8*: GACCCTAGGCAAGCTGTGG; CTGCACACGGAGCCTCTC

*Sox9*: TCGGACACGGAGAACACC; GCACACGGGGAACTTATCTT

*Dmrt1*: AAGGCCCCTCCTACTCAGAA; GCTGGAGAGGGAGACCAAG

*Gata1*: TGGGGACCTCAGAACCCTTG; GGCTGCATTTGGGGAAGTG

*Dhh*: CACGTATCGGTCAAAGCTGA; GTAGTTCCCTCAGCCCCTTC

*Ptgds*: GGCTCCTGGACACTACACCT; ATAGTTGGCCTCCACCACTG

*Hsd3b1*: GACCAGAAACCAAGGAGGAA; GCACTGGGCATCCAGAAT

*Cyp17a1*: CATCCCACACAAGGCTAACA; CAGTGCCCAGAGATTGATGA

*Cyp11a1*: AAGTATGGCCCCATTTACAGG; TGGGGTCCACGATGTAAACT

*StAR*: TTGGGCATACTCAACAACC; ACTTCGTCCCCGTTCTCC

*Srd5a1*: CATCTACAGGATCCCACAAGG; TCAATAATCTCGCCCAGGAA

*Trim28*: ATCAGCTGGCTACCGACTCT; GCACGAATCAAGGTCAGGTC

1*8S*: GATCCATTGGAGGGCAAGTCT; CCAAGATCCAACTACGAGCTTT

RNA-seq libraries were generated from 600 ng of total RNA using the TruSeq Stranded mRNA LT Sample Preparation Kit (Illumina, San Diego, CA), according to the manufacturer's instructions. Briefly, following purification with oligo d(T) magnetic beads, mRNA was fragmented using divalent cations at 94°C for 2 minutes. Cleaved RNA fragments were copied into first-strand cDNA using reverse transcriptase and random primers. Strand specificity was achieved by replacing dTTP with dUTP during the second-strand cDNA synthesis using DNA Polymerase I and RNase H. After addition of a single 'A' base and adapter ligation to double stranded cDNA fragments, products were purified and enriched by PCR (30 sec at 98°C; [10 sec at 98°C, 30 sec at 60°C, 30 sec at 72°C] x 12 cycles; 5 min at 72°C) to create the cDNA library. Surplus PCR primers were removed by purification with AMPure XP beads (Beckman-Coulter, Villepinte, France), and the quality and quantity of the final cDNA libraries were checked by capillary electrophoresis. Libraries were then sequenced on an Illumina HiSeq 4000 system using single-end 1 × 50 bp, following the Illumina recommendations. Image analysis and base calling were performed with RTA 2.7.3 and CASAVA 2.17.1.1. Reads were mapped to the mm10 assembly of the *Mus musculus* genome using STAR[90] version 2.5.3a. Gene expression quantification was performed from uniquely aligned reads using htseq-count[91] version 0.6.1p1, with annotations from Ensembl version 92 and "union" mode. Comparison between wild type and cKO samples was performed using the Wald test for differential expression proposed by Love et al.[84] and implemented in the Bioconductor package DESeq2 version 1.16.1.

**Testis and ovary dissociation for single cell RNA-seq analysis**

The testes from an 8-week-old control male were collected and enzymatically dissociated with a modified protocol from[92]. Briefly, tunica albuginea was delicately removed, and testes were incubated in DMEM (11885084; Gibco) supplemented with 1 mg/mL collagenase (C0130; Sigma-Aldrich), 1 mg/mL hyaluronidase (H3506; Sigma-Aldrich), and 0.8 mg/mL DNase I (dN25; Sigma-Aldrich) at

35°C for 10 minutes with gentle agitation. Tissue was centrifuged and incubated in DMEM (11885084; Gibco) supplemented with 1 mg/mL collagenase (C0130; Sigma-Aldrich), 0.025% trypsin-EDTA (25300-054; Gibco), and 0.8 mg/mL DNase I (dN25; Sigma-Aldrich) at 35°C for 25 minutes with gentle agitation. Cells were filtered through a 100 µm cell strainer and pre-stained with 100 µg Hoechst dye (B2261; Sigma-Aldrich) and 400 µg DNase I (dN25; Sigma-Aldrich) at 35°C for 20 minutes with gentle agitation. Trypsin was quenched with 600 µL foetal bovine serum (F2442; Sigma-Aldrich). Cells were stained with 6 µg Hoechst dye per million of cells (B2261; Sigma-Aldrich) and 40 µg DNase I (dN25; Sigma-Aldrich) at 35°C for 25 minutes with gentle agitation. Cells were centrifuged, resuspended in DMEM (11885084; Gibco) supplemented with 2% foetal bovine serum (F2442; Sigma-Aldrich) and filtered through a 70 µm cell strainer. Single cells were collected on a BD FACS Aria II by excluding debris (side scatter versus forward scatter), doublets (area versus width) and haploid cells with low DNA content (low Hoechst fluorescence intensity)[92]. FACS-sorting was performed at the Flow Cytometry Facility of the University of Geneva. Cells were centrifuged, resuspended in DMEM (11885084; Gibco) supplemented with 2% foetal bovine serum (F2442; Sigma-Aldrich), counted and immediately processed with a 10X Chromium controller.

Ovaries from two 8-week-old *Trim28*$^{fx/fx}$*; Nr5a1:Cre* females and two 8-week-old control females were collected, minced into small pieces and enzymatically dissociated in PBS (10010-015; Gibco) supplemented with 0.25 mg/mL collagenase (C0130; Sigma-Aldrich) and 0.0005% trypsin-EDTA (25300-054; Gibco) at 37°C for 30 minutes with gentle agitation. Trypsin was quenched with 200 µL PBS (10010-015; Gibco) supplemented with 3% BSA. Cells were centrifuged, resuspended in PBS (10010-015; Gibco) supplemented with 3% BSA and filtered through a 70 µm cell strainer. Single cells were collected on a BD FACS Aria II by excluding debris (side scatter versus forward scatter) and doublets (area versus width). FACS-sorting was performed at the Flow Cytometry Facility of the University of Geneva. Cells were centrifuged, resuspended in DMEM (11885084; Gibco) supplemented with 2% foetal bovine serum (F2442; Sigma-Aldrich), counted, and immediately processed with a 10X Chromium controller.

**Single cell RNA-seq library preparation, sequencing, and transcriptomic analyses**

Single-cell sequencing was performed on the 10X Genomic platform using 8-week-old *Trim28*$^{cKO}$ (*Trim28*$^{fx/fx}$*; Nr5a1:Cre* mouse line) (2 biological replicates) and control ovaries (2 biological replicates) and testes (1 biological replicate). To increase the numbers of testicular somatic cells, public 10X Genomic data from seven adult mice testes were incorporated[27] (Sup. Tab S2).

After converting the base calls to FASTQ format, reads were processed with the CellRanger v3 count module. This aligned reads with STAR[90] to the GRCm38 reference genome using the M15 (Ensembl 90) GENCODE annotation, and then derived the gene vs. cell expression counts matrices. These pre-processing steps were performed on the Baobab HPC cluster at the University of Geneva. The quality control statistics were checked, such as number of reads per cell and percentage of reads mapped to the reference transcriptome (Data S2). Using Scanpy with Anndata[93], matrices were concatenated across all samples, and cells with <50 genes expressed were removed (and genes expressed in <3 cells). This removed 1,683 of the 57,063 cells. Expression counts were normalized and log-transformed, and highly variably expressed genes were identified. The top 50 PCA components were embedded in the neighbourhood graph with batch correction via BBKNN[94] with the neighbours_within_batch=5 parameter that removed batch effects among biological replicates (fig. S6A, bottom right). Data were visualized in the transcriptional space with 2D uniform manifold approximation and projection (UMAP)[95] and force directed graphs (FDG) using the ForceAtlas2 implementation[96], mainly in Jupyter notebooks. Cell clusters were defined with the Leiden algorithm[97] and were annotated by marker gene analyses (fig. S6b). The supporting cell lineage was extracted, represented by Leiden clusters 2 (granulosa/mutant) and 35 (Sertoli). Cluster 2 expressed the granulosa cell markers *serpine2*, *Hsd17b1*, *Inhba*, *Amh*, *Foxl2*, *Nr5a2*, *Fst*, *Wnt4*, *Esr2*. Cluster 35 expressed the Sertoli cell markers *Cldn11*,

*Sox9*, *Sox8*, *Rhox8*, *Ptgds*, *Gata1*, *Nr0b1*, *Abtb2*. Then, the Leiden clustering was repeated at a higher resolution to separate the granulosa, intermediate, and Sertoli cell populations in three distinct populations that were compared by partition-based graph abstraction (PAGA) (Fig. 2a, right)[98]. These cells were ordered along a diffusion pseudo-time[31], setting the starting root cell in the granulosa control cells, and were plotted according to the expression of the differentially expressed genes calculated with Wilcoxon rank-sum tests (with Benjamini-Hochberg multiple testing correction) among the three clusters, filtering genes that had a fold change <1.5 and genes that were expressed in ≥0.8 proportion of the cluster in which they were downregulated. The top 70 upregulated genes were plotted for each cluster in Fig. 2d, and the full sets of filtered upregulated genes is shown in Sup. Tab S3. The mean expression across marker genes was also plotted using previously defined modules: F23 for granulosa, F17 for Sertoli, and F1-F10 for progenitor cell markers; see Figure 5 in[30]. The functional annotations that were enriched among gene lists were examined using hypergeometic tests performed via g:Profiler[99]. To calculate the enrichments in Sup. Tab. S4, 70 upregulated genes were used in the intermediate cell population as the foreground gene set, and all the mouse genes expressed in the concatenated Anndata matrix after pre-processing as the background gene set.

**Antibodies**

Polyclonal antibodies against SUMO-1 and SUMO-2/3 were produced by injecting rabbits with recombinant His-tagged mouse SUMO-1 and SUMO-3 proteins produced in bacteria and purified using Ni-NTA column followed by Superdex 75 gel filtration, as previously described[100]. Rabbit sera were affinity-purified using GST-SUMO1 or GST-SUMO2 coupled to CnBr-activated Sepharose (SIGMA). Their specificity was confirmed by immunoblotting using recombinant mouse SUMO-1 and SUMO-2. Dilution for immunofluorescence: 1/300 For ChIPseq: 2µg /IP

Other antibodies:

FOXL2 (rabbit)[101]. Immunofluorescence: 1/400. ChIP-seq and ChIP-SICAB: 5µl (2µg) / IP.

SOX9 (rabbit)[102]. Immunofluorescence : 1/400.

SOX8 (guinea pig)[103]. Immunofluorescence : 1/300.

DMRT1 (rabbit)[104]. Immunofluorescence: 1/400.

TRIM28 (mouse)[105]. Immunofluorescence: 1/1000.

TRIM28 (rabbit)[106]. ChIP-seq: 2µg / IP.

V5-HRP: Invitrogen. Ref P/F 46-0708. Western blotting:1/5000

HA-HRP:. Invitrogen. Ref 26183-HRP. Western blotting: 1/5000

V5: Invitrogen. Ref 37-7500. Western blotting: 1/2000.

HA: Invitrogen. Ref 21183. Western blotting: 1/2000.

FLAG: Invitrogen. Ref MA1-91878. Western blotting: 1/3000

Tubulin: Sigma. Ref T9026. Western blotting: 1/3000

**Histology, immunofluorescence, and confocal microscopy**

Post-natal and adult ovaries were collected, fixed in 4% paraformaldehyde, and paraffin-embedded. Then, 4-µm sections were cut and processed for histology and immunofluorescence. Sections were stained with periodic acid-Schiff (PAS) stain using standard protocols. Immunofluorescence was performed as previously described[107]. Overnight incubation with primary antibodies at 4°C was followed by incubation with the appropriate secondary antibodies (1/800) (Alexa-Ig, Molecular Probe). Nuclei were stained with DAPI (Sigma). Images were captured with a Zeiss Axioimager Apotome microscope. For quantification of SUMO1 and SUMO2 immunofluorescence signal, images were captured with a Zeiss LSM780 confocal microscope and analysed with the Imaris. Statistical analyses were performed using the GraphPad Prism v9 software using two-way ANOVA with the Dunnett's multiple comparisons test.

The Flag-TRIM28 expression plasmid was described previously[105]. The TRIM28 C651F mutation was generated using the Quick-change II site-directed PCR mutagenesis kit (Agilent) and the sequence was verified by Sanger-sequencing. The pcDNA3-HA-SUMO2 expression plasmid was obtained from Dr Ronald Hays. The pcDNA3.1-FOXL2-V5 (V5-tagged in C-terminal) expression plasmid was obtained from Prof Reiner Veitia. To generate the pcDNA3.1-RUNX1-V5 construct (i.e., RUNX1 C-terminally tagged with V5), the human RUNX1 open reading frame was PCR-amplified from the RUNX1-pCSdest plasmid (a gift from Roger Reeves. Addgene plasmid # 53802; http://n2t.net/addgene:53802; RRID:Addgene_53802) and subcloned in the pcDNA3.1/V5-HIS TOPO vector (Invitrogen) following the manufacturer's instructions. The sequence was verified by Sanger-sequencing.

For SUMOylation assays in transfected cells: HEK293T cell culture, transfection, and SUMOylated FOXL2 and RUNX1 immunoprecipitation conditions were as described previously[53]. FOXL2-V5 and RUNX1-V5 were immunoprecipitated using V5 agarose affinity gel (SIGMA ref: A7345). IP complexes were analysed by western blotting using anti-V5 and anti-HA HRP-coupled antibodies (see Antibodies section). For inputs, crude cellular extracts were probed with non-coupled antibodies (see Antibodies section).

**Steroid quantification**

Steroids (testosterone, androstenedione, β-oestradiol and estrone) were quantified in gonads from 4-month-old control females, *Trim28*$^{cKO}$ females, and control males (n=4/group) by liquid chromatography coupled to tandem mass spectrometry (LC-MS/MS). Briefly, adult tissues were homogenized in methanol containing mixed internal standards. After evaporation, dried residues were resuspended in 600 µl of methanol, loaded onto an Oasis Prime HLB (Waters) cartridge, and sequentially washed with 0.1% formic acid in 35% methanol, and 0.1% ammonia solution in 35% methanol. Samples were eluted with 45 ml of methanol and then diluted with 55 µl of distilled water.

After mixing, 30 µl of each sample was injected for Ultra High-Performance Liquid Chromatography (UPLC) analysis. LC-MS/MS analysis was performed on a UPLC Acquity system (Waters Corporation) coupled to a triple-quadrupole XevoTQD mass spectrometer (Waters Corporation). Steroids were analysed with a reversed-phase column (Acquity UPLC HSS T3 column C18, 1.8 µm, 2.1x50mm, Waters Corporation). The chromatographic mobile phase was 2 mM ammonium acetate with 0.1% formic acid (v/v) in water (Phase A) and 2 mM ammonium acetate with 0.1% formic acid (v/v) in methanol (Phase B) delivered at a flow rate of 0.6 mL/min at 50°C. The gradient was 0-1 min, 55% A; 1-3.5 min, 50% to 35% A; 3.5-3.51 min, 35% to 5% A. To equilibrate the column again, at 3.51 min the gradient remained in the initial conditions for 1.49 min. The total run time was 5 min. Steroid ionization was performed using positive electrospray ionization of ($[M+H]^+$) with the following settings: capillary voltage, 500 V; cone voltage, 35 V; desolvation gas, 1000 L/h; cone gas, 80 L/h; desolvation temperature, 500 °C; and source temperature, 150 °C.[108]

**Data and material availability**

All data are available in the main text or supplementary materials. RNA-seq and ChIP-seq data have been deposited in the Gene Expression Omnibus under accession number GSE166385 (RNA-seq and ChIP-seq) and GSE166444 (scRNA-seq) and the mass spectrometry proteomic data have been deposited in the ProteomeXchange Consortium via the PRIDE partner repository with the dataset identifier PXD024439.

**Acknowledgments**

This study was supported by the Centre National de la Recherche Scientifique (CNRS), the University of Montpellier, and by a grant from 'Agence Nationale de la Recherche' (ANR-16-CE14-0020-SexMaintain to F.P). R.M and R.L.B are funded by the Francis Crick Institute. The Francis Crick

Institute receives its core funding from Cancer Research UK (FC001107), the UK Medical Research Council (FC001107), the Wellcome (FC001107), and by the UK Medical Research Council (U117512772). We thank Nitzan Gonen from Bar-Ilan University for scientific discussions and critical reading of the manuscript. We are grateful to Monsef Benkirane and Dominique Giorgi for their continual support to the project. We thank the staff of the 'Reseau d'Histologie de Montpellier'(RHEM) for paraffin embedding of tissue samples and histology staining. We thank Marie-Pierre Blanchard, and Amelie Sarrazin of the Montpellier Imaging Facility (MRI) for their help with microscopy experiments. We thank the staff of the RAM-CECEMA animal care facility (University of Montpellier). We thank Francoise Kuhne for technical assistance in single-cell analysis. We thank the GenomEast platform, particularly Violaine Alunni for the preparation of RNA-seq libraries and Romain Kaiser for sequencing. We thank Prof Michael Wegner and Prof David Zarkower for their generous gift of the anti-SOX8 and anti-Dmrt1 antibodies respectively. We thanks Prof Reiner Veitia for the generous gift of the pcDNA3.1-FOXL2-V5 expression plasmid. We thank the Freiburg Galaxy server that was used for some calculations. We thank Elisabetta Andermarcher for manuscript editing.

**Author contribution**

M.R and F.P designed the study. M.R, S.D, L.L and B.B.B performed histological and RT-qPCR experiments. S.N and F.P designed the scRNA-seq. C.R, MR, Y.N and S.N. performed and analyzed scRNA-seq. F.P designed and performed ChIPseq experiments. All bioinformatics analyses, excepted for scRNA-seq, were performed by S.L. F.C designed and created the mouse line carrying $Trim28^{Phd}$ allele. A.P and A.L.N performed steroids dosage. G.B and D.W produced and purified the SUMOs and FOXL2 antibodies respectively. R.L.B, R.M and M.R.R designed, performed and analysed FOXL2-ChIP-SICAP. F.P wrote the manuscript. All authors reviewed and added input to the manuscript.

### Competing interests

The authors declare no competing interests.

### Correspondence and requests for materials

should be addressed to F.P

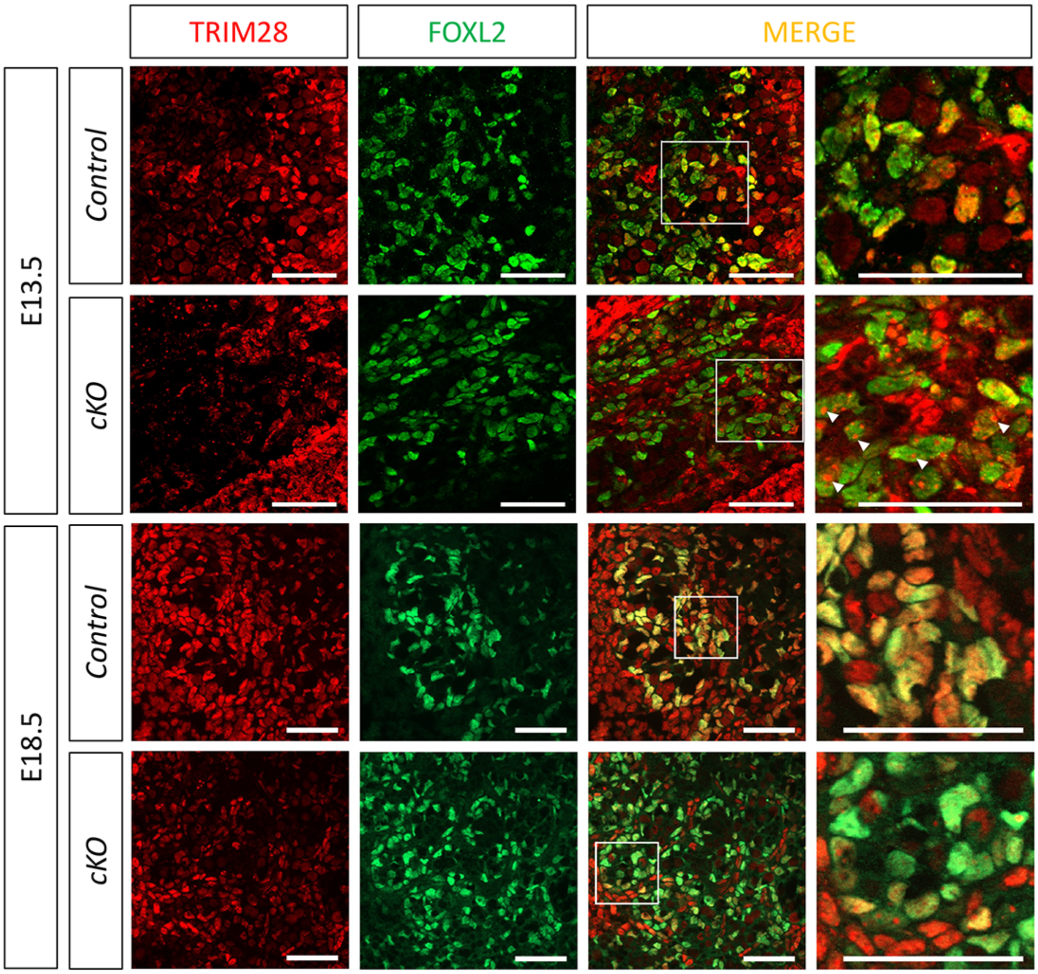


**Fig. S1**. Expression of the TRIM28 protein in foetal pre-granulosa cells of control and mutant ovaries. At 13.5 dpc (E13.5) and 18.5 dpc (E18.5) in XX gonads, TRIM28 (red) is co-expressed with FOXL2 (green) in the nucleus of pre-granulosa cells. At E13.5, TRIM28 is in nucleoplasm and concentrated in nuclear dots that might be heterochromatin. At E18.5 TRIM28 is diffusely located within the nucleoplasm. In mutants, at E135, TRIM28 is in nuclear dots (white arrowheads) and appears decreased in the nucleoplasm of pre-granulosa cells. In E18.5 XX mutants, TRIM28 has nearly disappeared from the nucleus of pre-granulosa cells expressing FOXL2. Scale bar: 20µm.

**Fig. S2.** Upper panels: PAS staining showing that at 3 days post-partum (3 dpp), *Trim28*$^{cKO}$ (*cKO*) ovaries display the same structures observed in control ovaries. Scale bar: 100µm. Lower panels: Double immunofluorescent staining shows that TRIM28 and FOXL2 are co-expressed in immature granulosa cells of 3 dpp control ovaries. In 3 dpp *Trim28*$^{cKO}$ ovaries, TRIM28 signal has almost disappeared from cells that express FOXL2. Red staining in *Trim28*$^{cKO}$ ovary sections corresponds to oocytes where *Trim28* is not deleted. Double staining for SOX8 and SOX9 shows that these Sertoli cell markers are not expressed in granulosa cells from 3 dpp control and *Trim28*$^{cKO}$ ovaries (Ov). Conversely, in 3 dpp testes (Control Test), SOX8 and SOX9 are co-expressed in Sertoli cells that form seminiferous tubules. Interstitial staining corresponds to a secondary antibody artifact. Scale bar: 50µm.

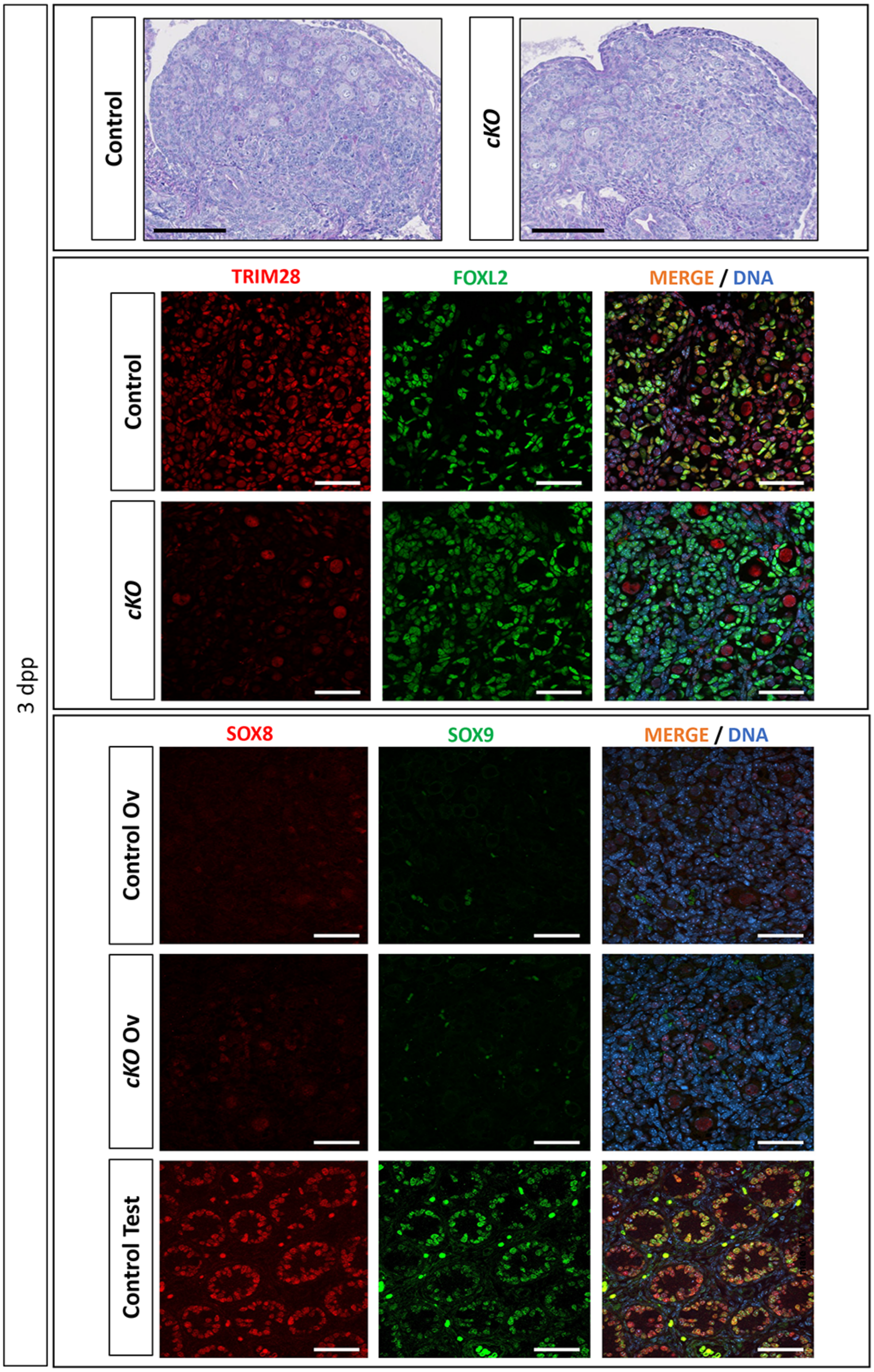

**Fig. S3.** Double staining for FOXL2 and SOX8, FOXL2 and SOX9, and SOX8 and SOX9 in week 8 post-partum *Trim28*$^{cKO}$ ovaries. Transdifferentiation of supporting cells is more advanced in the medullar (Med) than in the cortical area (Cor) (arbitrarily separated by a dotted white line). Boxed areas in the cortical and medullar areas are shown at higher magnification. Ovarian cortex still displays organized follicular structures that express FOXL2, although some isolated cells express SOX8 and/or SOX9. Conversely, in medullar areas, most follicular structures have disappeared and are reorganized in pseudo-tubules that express SOX8 and SOX9. SOX8 and SOX9 signals frequently overlap, but rarely with FOXL2. Strong interstitial staining is due to secondary antibody artifacts. Scale bar: 50µm.

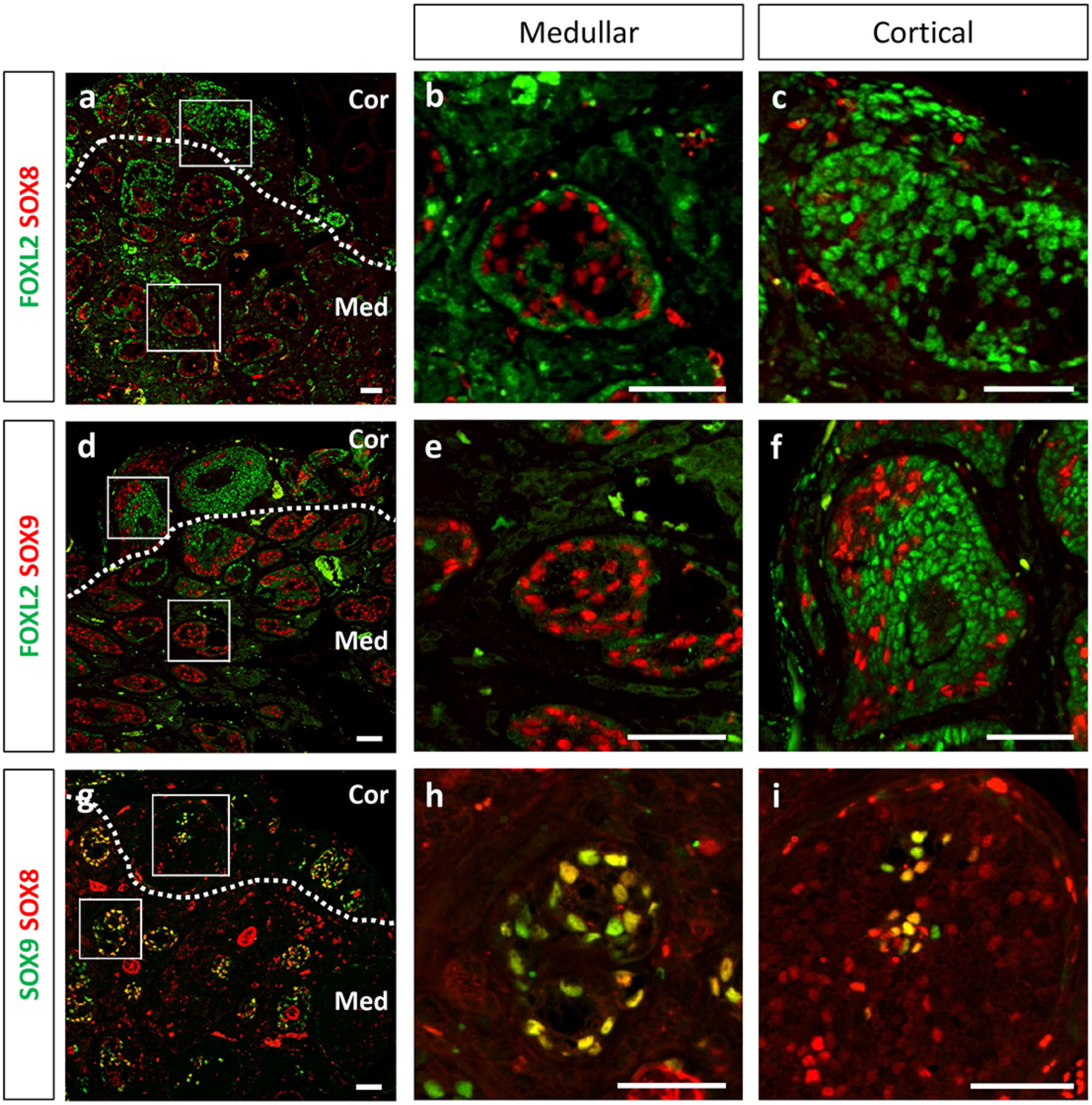

**Fig S4.** TUNEL staining of XX control and *Trim28*$^{cKO}$ ovaries at 20 dpp and 8 weeks showing apoptotic nuclei (dark brown). No obvious difference was observed between follicles from control and mutant ovaries. Scale bar 50 μm.

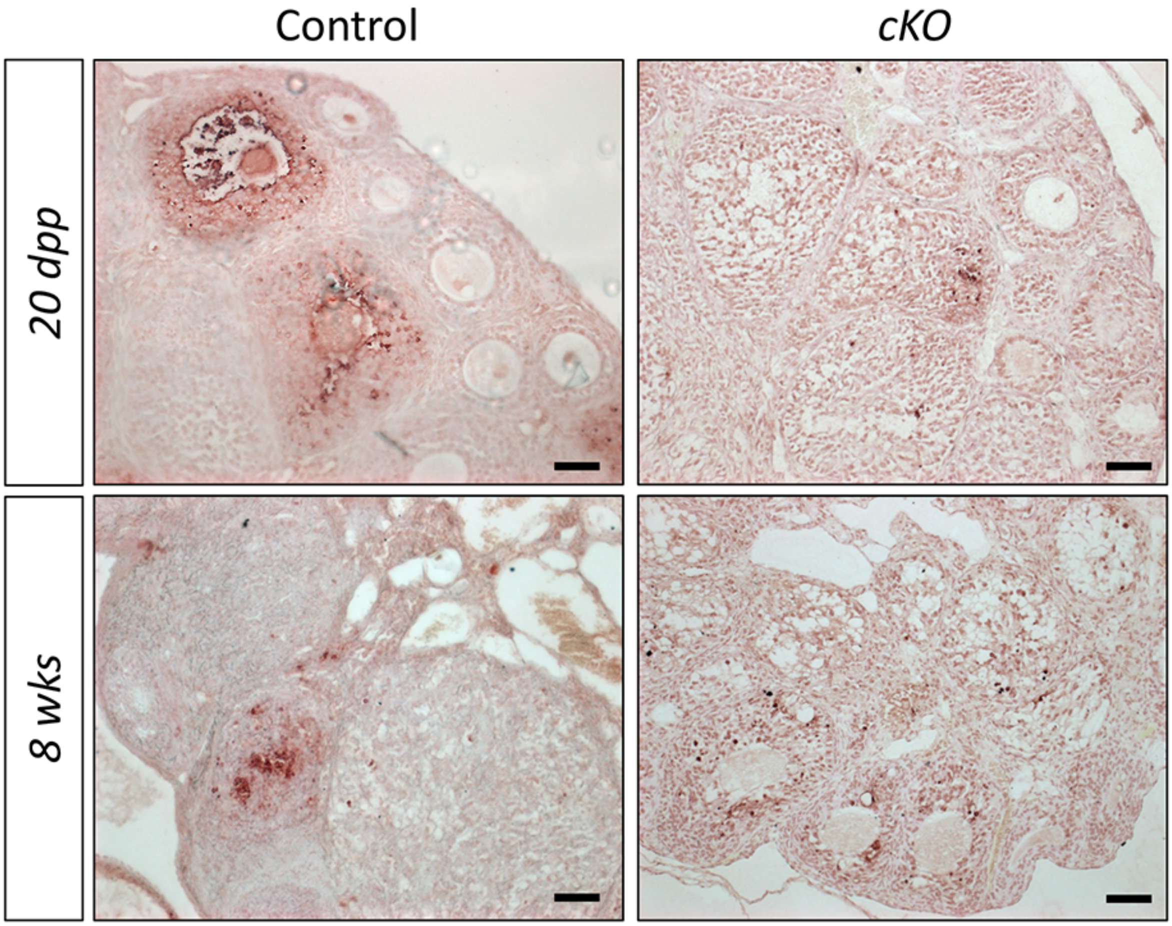

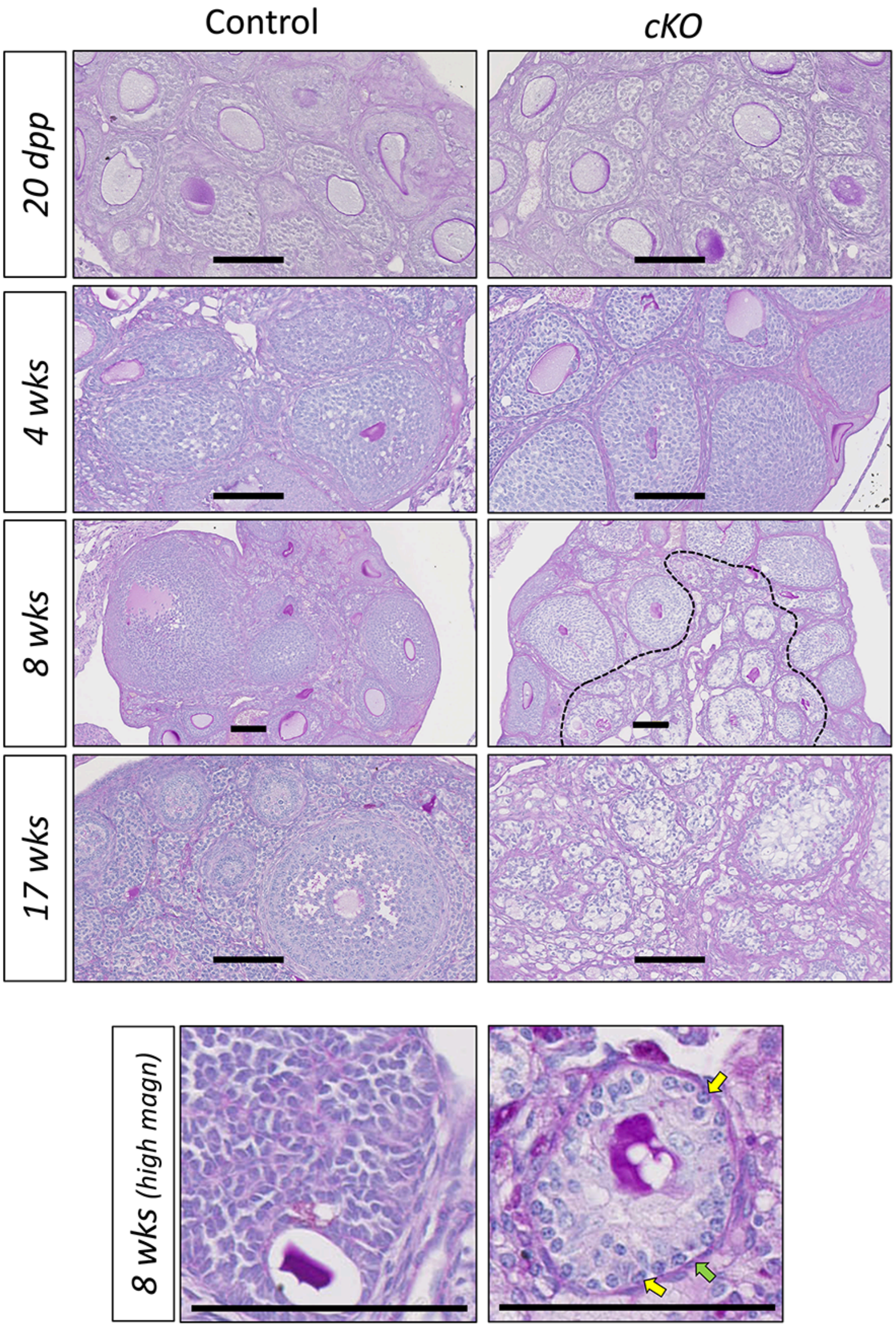


**Fig. S5.** Progressive reorganization of ovarian follicles in *Trim28*$^{cKO}$ (*cKO*) ovaries from 20 dpp to week 17 post-partum, visualized by PAS staining. In control, ovarian follicles develop normally, from pre-antral to pre-ovulatory follicles. At week 4 post-partum, ovarian organization is similar in control and *cKO* animals. At week 8 post-partum, multiple follicles reorganized in pseudo-tubules delineated by basal laminae are observed in the medullar region of *cKO* ovaries (arbitrarily delimited by a dotted black line). Cortical follicles contain apparently normal oocytes, while in medullar pseudo-tubules oocytes are degenerating or have disappeared. At week 17 post-partum, in *cKO* ovaries supporting cell transdifferentiation has spread to the entire ovary, and no oocyte can be detected. Bottom, High magnification of 8-week-old ovary tissue sections showing a preantral follicle and pseudo-tubules with features of Sertoli cells: nuclei with tripartite nucleoli (yellow arrows) and important deposition of basal laminae (green arrow). Scale bar: 100µm.

**Fig. S6.** Effect of *Trim28* deletion heterozygosity on post-natal ovary. **a.** Double immunofluorescent staining for TRIM28 (red) and FOXL2 (green) in adult ovaries at 20 dpp, 4 weeks, and 8 weeks. No visible difference between wild type (Trim28$^{+/+}$; Nr5a1:Cre) and heterozygous mutant (Trim28$^{fl/+}$; Nr5a1:Cre) ovaries was observed in FOXL2 staining unlike in the homozygous mutant (Trim28$^{fl/fl}$; Nr5a1:Cre). Scale bar: 50µm.
**b.** RT-qPCR analysis of the Sertoli cell markers *Sox9*, *Sox8* and *Dmrt1* in 3-month-old ovaries from wild type (+/+: Trim28$^{+/+}$; Nr5a1: Cre), heterozygous mutant (fl/+: Trim28$^{fl/+}$; Nr5a1:Cre), and homozygous (fl/fl: Trim28$^{fl/fl}$; Nr5a1:Cre) mutant mice. Bars are the mean ±SEM, *n* is indicated for each condition. ****$P$ <0.0001, **$P$<0.05, (one-way ANOVA with Tukey's multiple comparisons test).

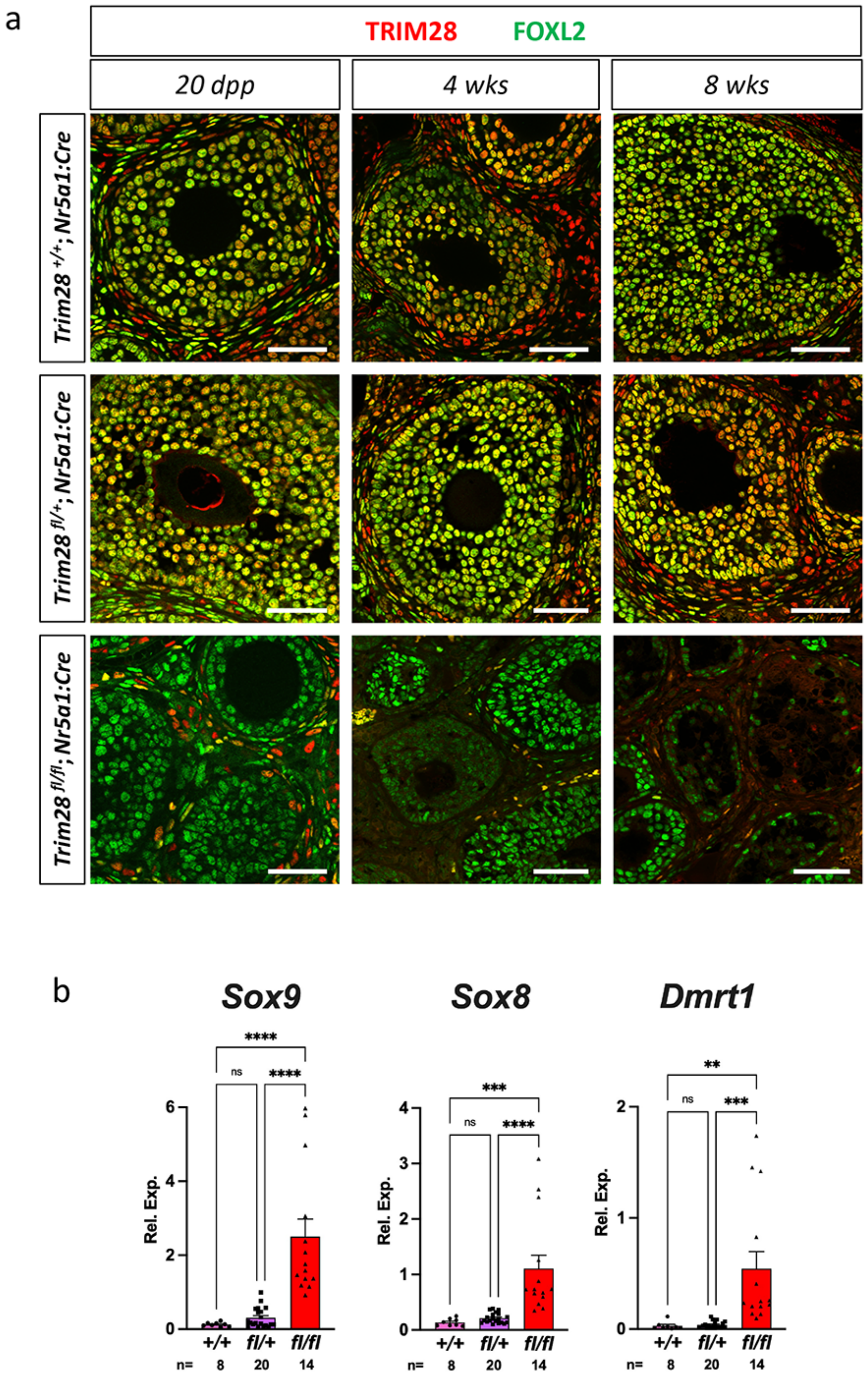

**Fig. S7.** The expression profile of several genes involved in steroidogenesis is modified in *Trim28*$^{cKO}$ ovaries. **a,** RT-qPCR analysis of key steroidogenesis genes from 0.5 to 4 months post-partum in control (Control Ov) and *Trim28*$^{cKO}$ (cKO) ovaries, and control testes (Cont test). Statistical data are provided in Source data file. **b,** Expression of genes involved in steroidogenesis assessed in 7-month-old ovaries by RNA-seq (n=3). Values correspond to normalized read counts divided by the median of the transcript length in kb. Adj P Val<0.05. │Log2FC│>1. Values are the mean ± SD.

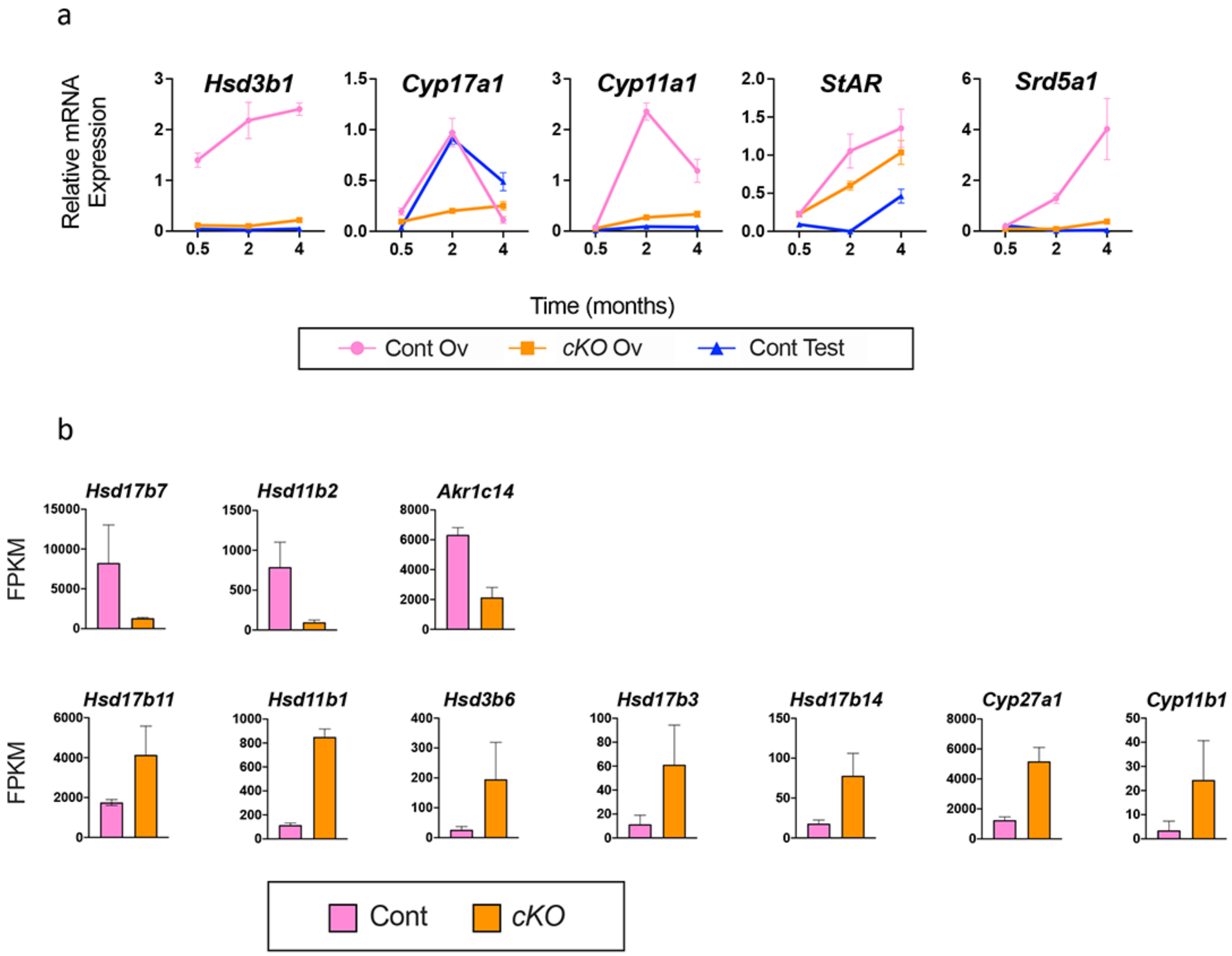

**Fig. S8.** Single-cell atlas of 8-week-old *Trim28*$^{cKO}$ and control ovaries and testes. **a,** UMAP representations of single cells in the transcriptional space coloured according to the Leiden clustering (left), condition (i.e. ovary/testis and mutant/control; upper right) and biological replicate (lower right). **b,** Expression of marker genes in the different Leiden clusters (1 to 42). Details in Data S3, tab df_gene_clusters). The colour of each dot indicates the mean expression within that cluster based on the normalized plus log transformed counts. The dot size represents the fraction of cells expressing that gene. Note that not all markers are specific, for example *Star* is a steroidogenic marker of both Leydig and theca cells. Cells from clusters 2 and 35 are granulosa/mutant and Sertoli cells, respectively, and were used in the subsequent analyses.

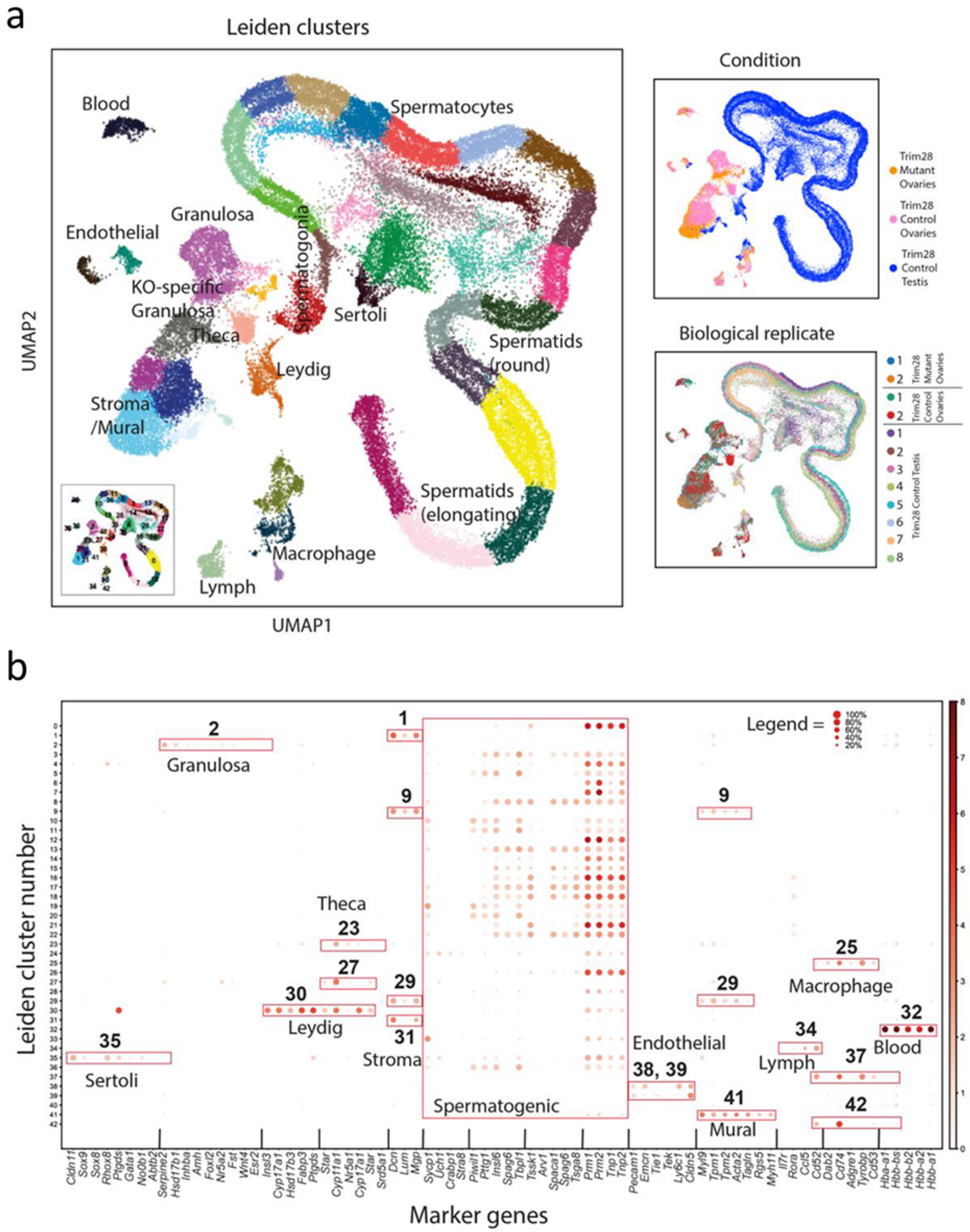

**Fig. S9.** *Trim28* expression is decreased in the cKO mutant compared with control ovaries. Violin plots show the expression of *Trim28* in all ovarian cell types (**a**) and only in supporting cells (**b**). The Wilcoxon rank sum test across all genes to calculate the multiple-testing adjusted p-values and log fold-changes demonstrated that *Trim28* expression levels are lower in mutant than control ovaries.

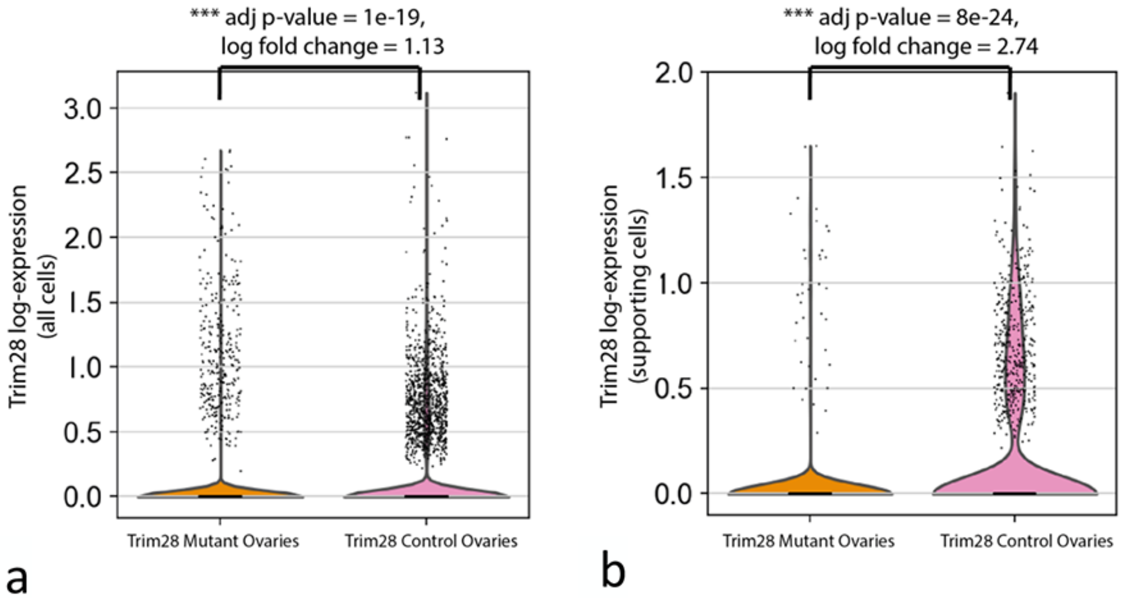

**Fig. S10**. Expression of granulosa and Sertoli marker genes in supporting cells (i.e. granulosa, intermediate, and Sertoli) across the binned pseudo-time.
**a,** Representation of the binary (yes/no) expression of *Sox9*, *Sox8* and *Dmrt1* on the force directed graphs (FDG) from **Figure 1**.
**b,**. Ten different pseudo-time bins are displayed on the FDG representing the pseudo-time value as a percentage, p, as follows: 0≤p≤10, 10<p≤20, 20<p≤30, 30<p≤40, 40<p≤50, 50<p≤60, 60<p≤70, 70<p≤80. 80<p≤90, 90<p≤100. Violin plots showing the expression of selected granulosa (**c**) and Sertoli (**d**) cell markers across the binned pseudo-time.

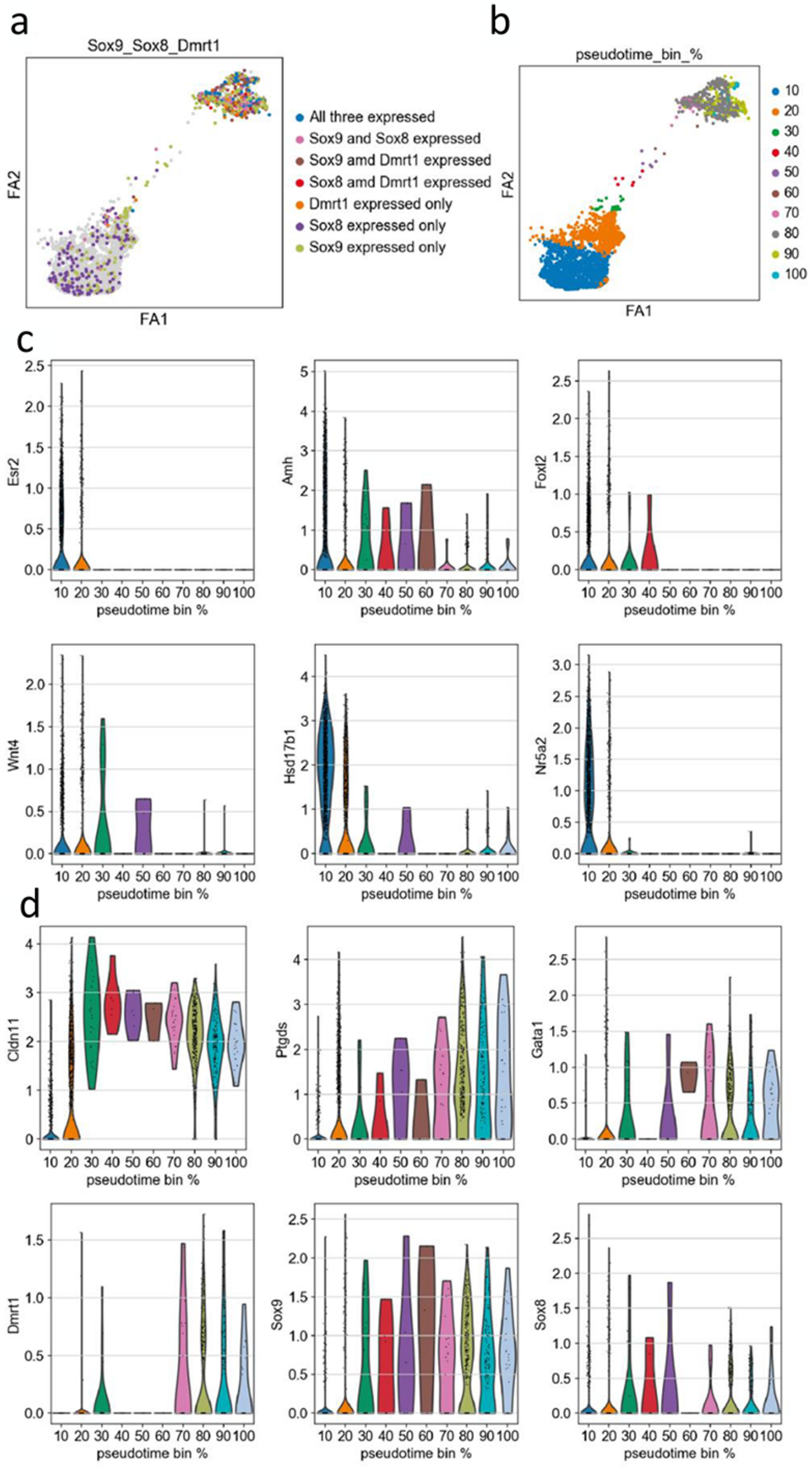

**Fig. S11**. Distribution of TRIM28 and FOXL2 ChIP-seq peaks on genes expressed in granulosa cells that are downregulated in *Trim28*$^{cKO}$ ovaries. Relevant ChIP-seq peaks are highlighted by a blue (TRIM28) or red (FOXL2) background.

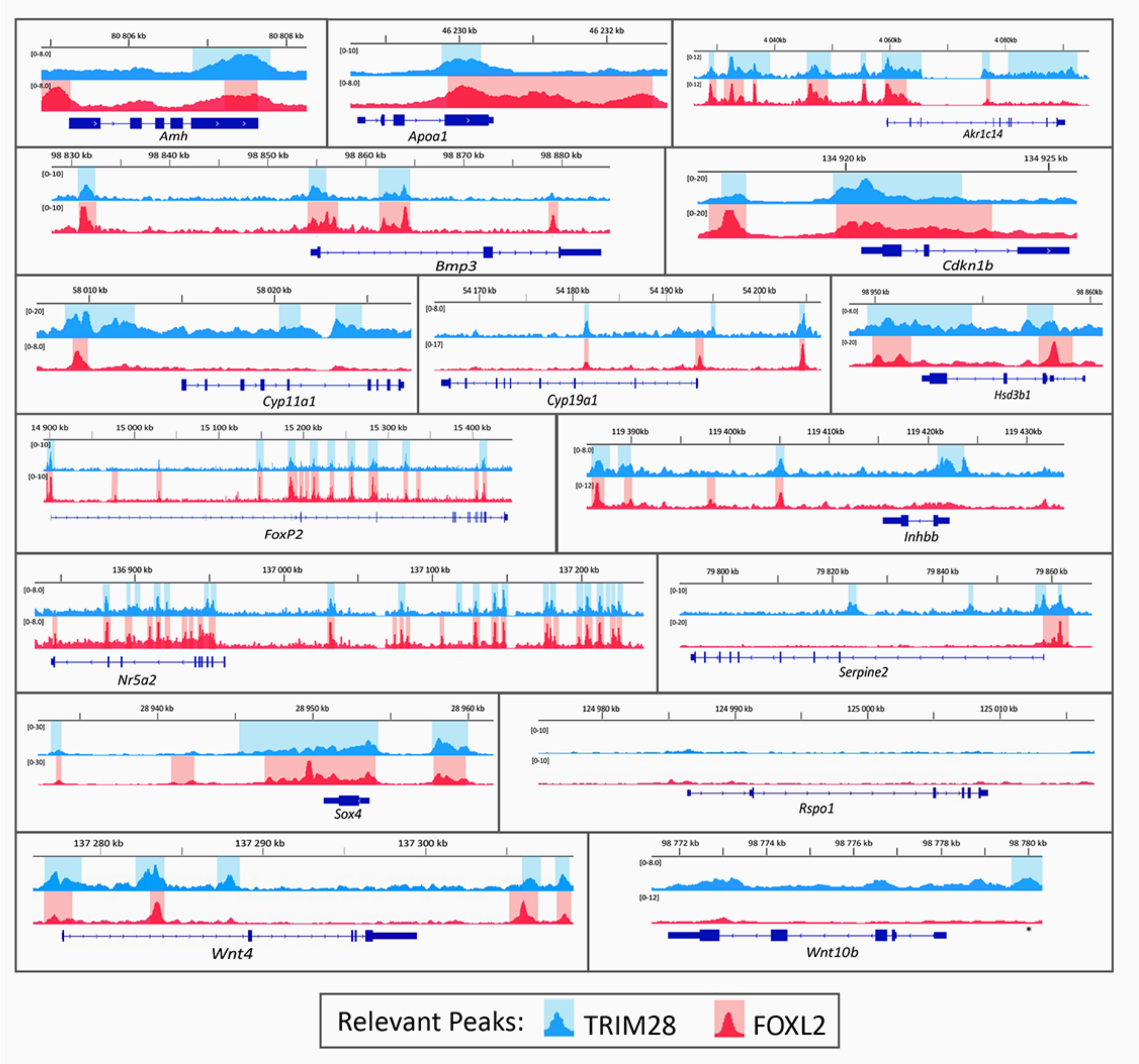

**Fig. S12**. Same as in fig. S11 but for genes expressed in Sertoli cells that are upregulated in *Trim28*$^{cKO}$ ovaries. Relevant ChIP-seq peaks are highlighted by a blue (TRIM28) or red (FOXL2) background.

.

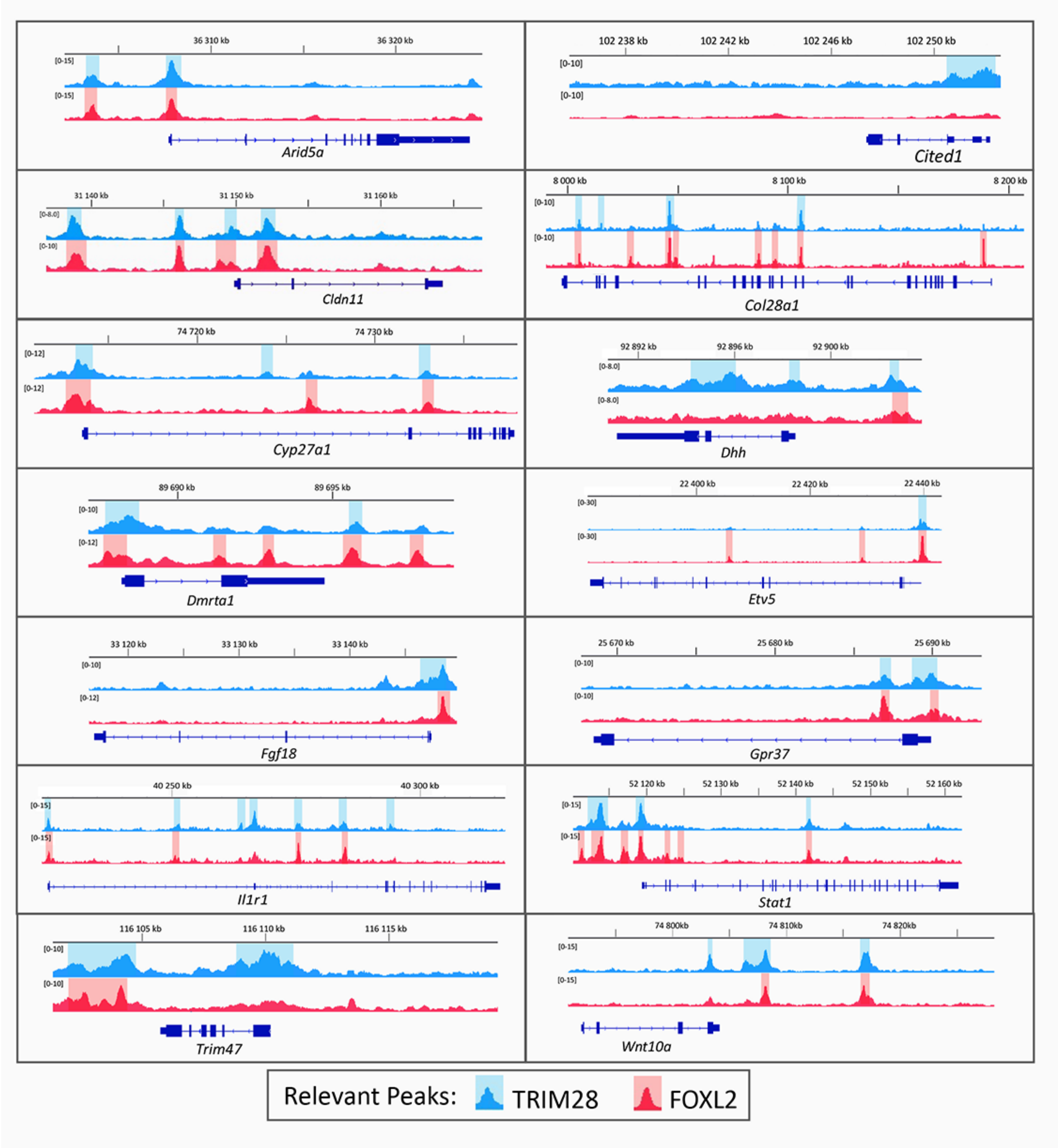

**Fig. S13.** Targeted mutation (C651F) of the PHD domain of TRIM28. Diagram showing the genomic map of the mouse *Trim28* gene; the targeting constructs to introduce the C651F mutation in the PHD domain (exon 13); Cre-mediated excision of the loxP-site-flanked sequences. Exons are represented as black boxes and introns as connected lines. The loxP sites are represented by red triangles and the PGK-neo cassette is indicated.

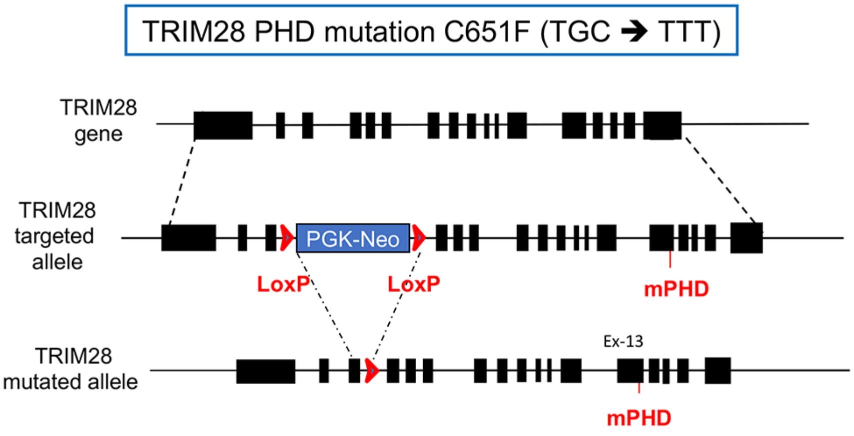


**Fig. S14.** Immunofluorescent staining for TRIM28 in *Trim28*$^{+/+}$ control and *Trim28*$^{Phd/cKO}$ ovaries showing that the mutant TRIM28$^{C651F}$ protein is effectively produced and localizes in the nucleus. Boxed areas are shown at higher magnification (lower panels) Compared with control, high magnification view of TRIM28$^{C651F}$ *(PHD/cKO)* shows a lower nuclear staining because only the mutated allele is expressed. Scale bar: 10µm.

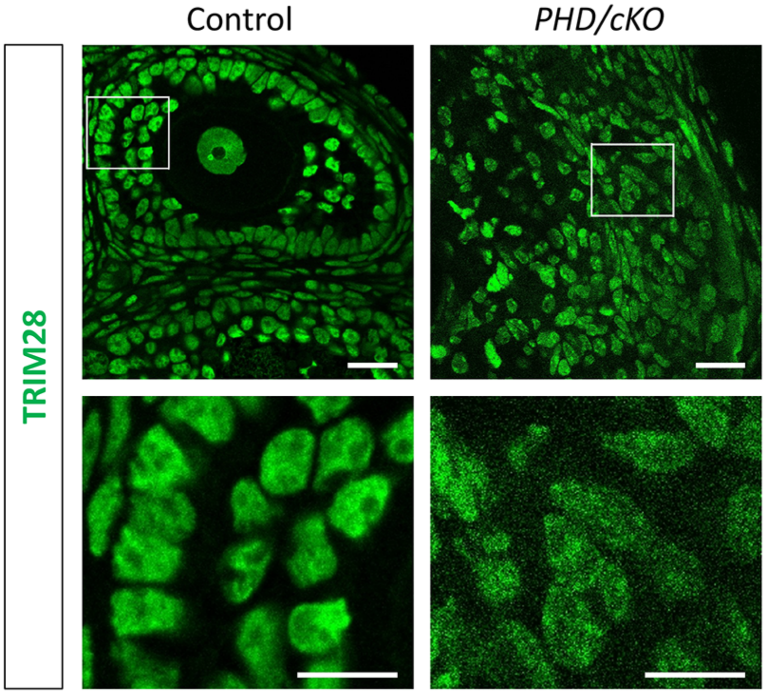

**Fig. S15.** PAS staining of medullar regions of 8-week-old ovaries showing normal follicular structures in *Trim28*$^{Phd/+}$ *(PHD/+)* and in control (*Cont*) ovaries. Conversely, *Trim28*$^{Phd/cKO}$ (*PHD/cKO*) ovaries display disorganized follicles with appearance of pseudo-tubules, as observed in *Trim28*$^{cKO}$ (*cKO*) ovaries. Scale bar 100µm.

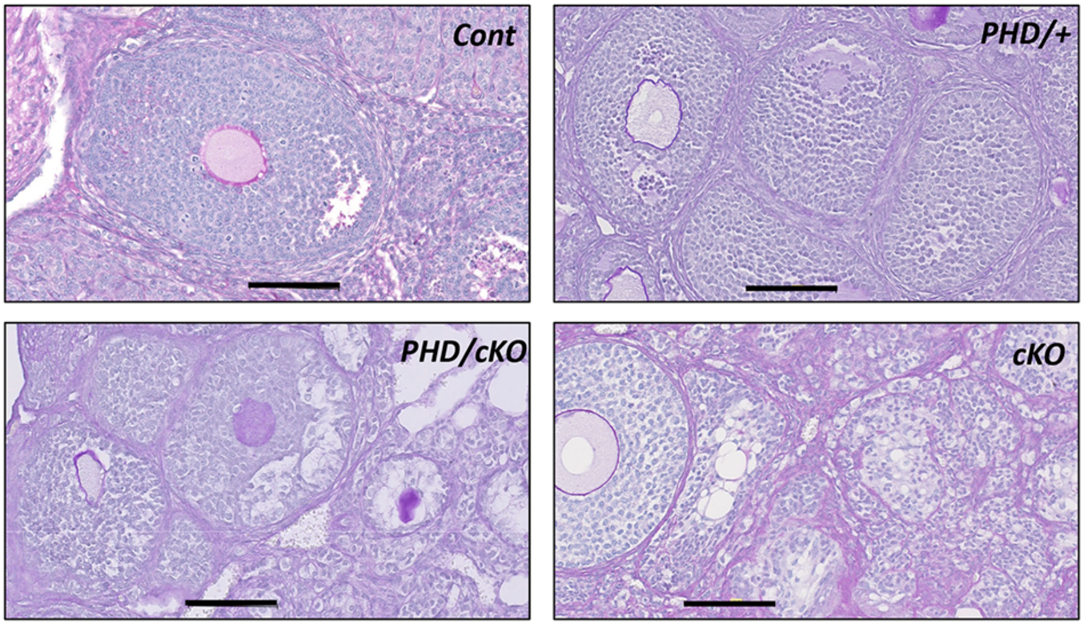


**Fig. S16.** MA plots for SUMO1 (**a** and **b**) and SUMO2 (**c** and **d**) showing the mean *versus* the ratio of normalized read counts. Spots corresponding to differential signal between mutant (*Trim28*$^{cKO}$, *cKO,* or *Trim28*$^{Phd/cKO}$, *PHD*) and control with Adj Pval<0.05 coloured in blue (hypo-SUMOylated) or red (hyper-SUMOylated).

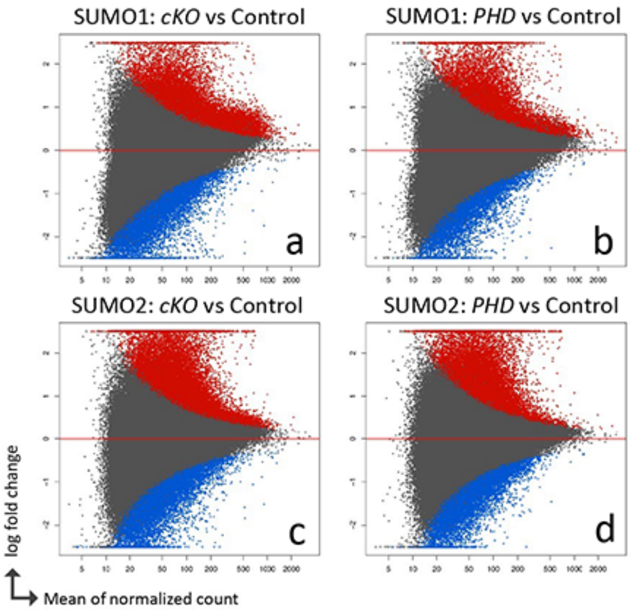

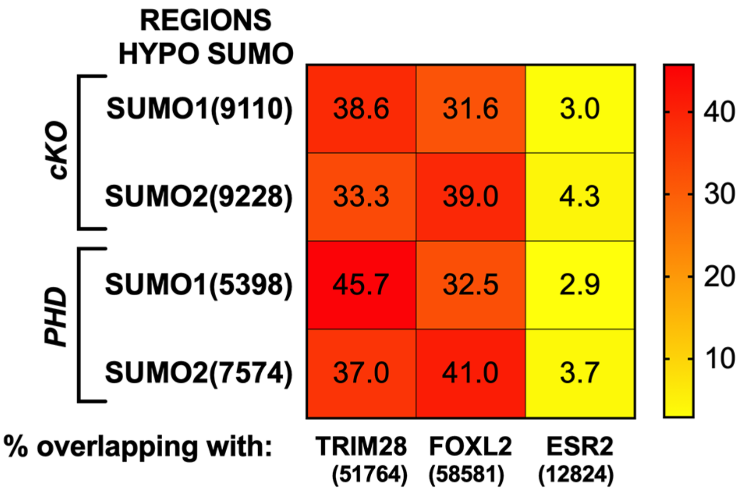


**Fig. S17.** Percentage of hypo-SUMOylated ChIP-seq peaks (SUMO1 and SUMO2) in mutant ovaries (*Trim28*$^{cKO}$*, cKO,* and *Trim28*$^{Phd/cKO}$, *PHD*) that overlap with TRIM28 and FOXL2 peaks genome-wide in control ovaries. ESR2 peaks where obtained from Lindeman and colleagues[60] The values in brackets correspond to the number of peaks.

**Fig. S18:** TRIM28 induces SUMOylation of FOXL2 and RUNX1. HEK293T cells were transfected with plasmids expressing V5-tagged (C-terminal) FOXL2 or RUNX1, HA-SUMO2, and Flag-tagged wild type TRIM28 or TRIM28 C651F. After 48h of transfection, cells were lysed in buffer with high concentration of SDS, diluted, and then immunoprecipitated with anti-V5 affinity resins. The IP complexes were analysed by western blotting with an anti-HA antibody to detect SUMOylated proteins and an anti-V5 antibody (upper panels: IP-V5). The IP complexes and cell lysates were also probed with the indicated antibodies to determine the protein input and the overall SUMOylation of cellular proteins (lower panels: Input). For both FOXL2 and RUNX1, the SUMOylated proteins of lower molecular weight correspond to the band detected by the V5 antibody suggesting that both proteins are SUMOylated by an endogenous E3-ligase. By contrast, in the presence of TRIM28, a SUMOylated form of higher molecular weight is detected for FOXL2 and RUNX1 (red triangle), but not in cells transfected with TRIM28 C651F.

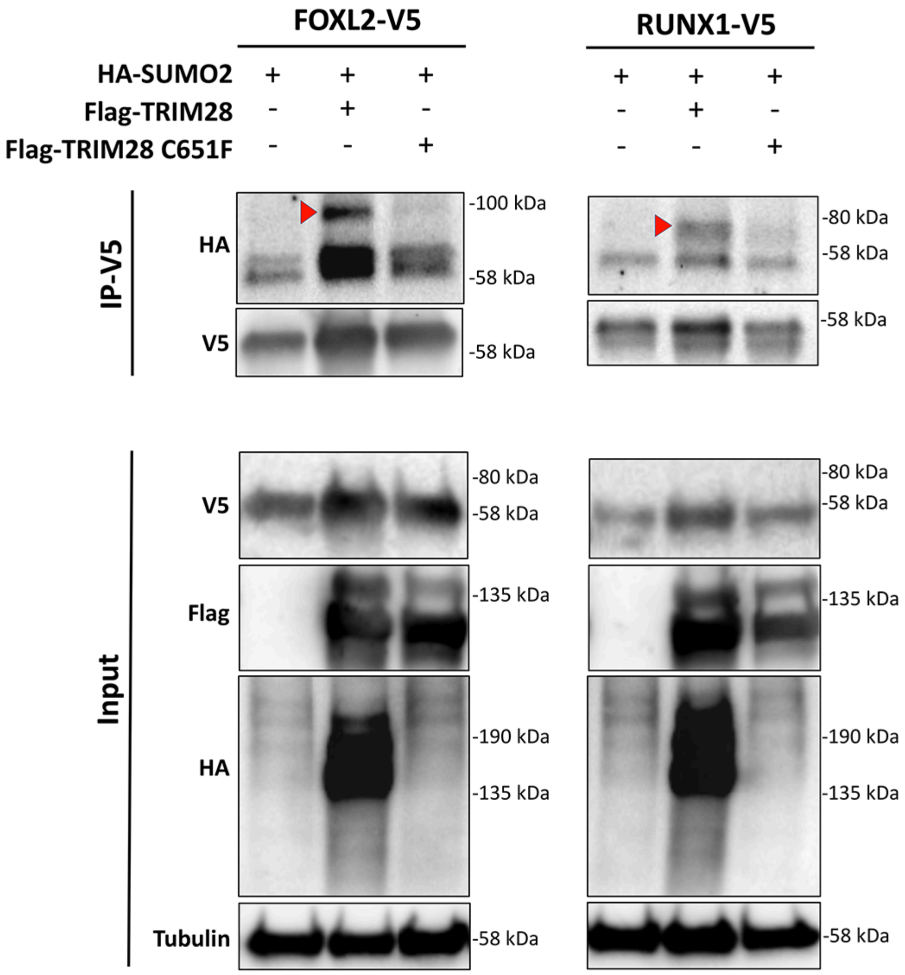

**Fig S19.** Percentage of overlap between the indicated peaks.

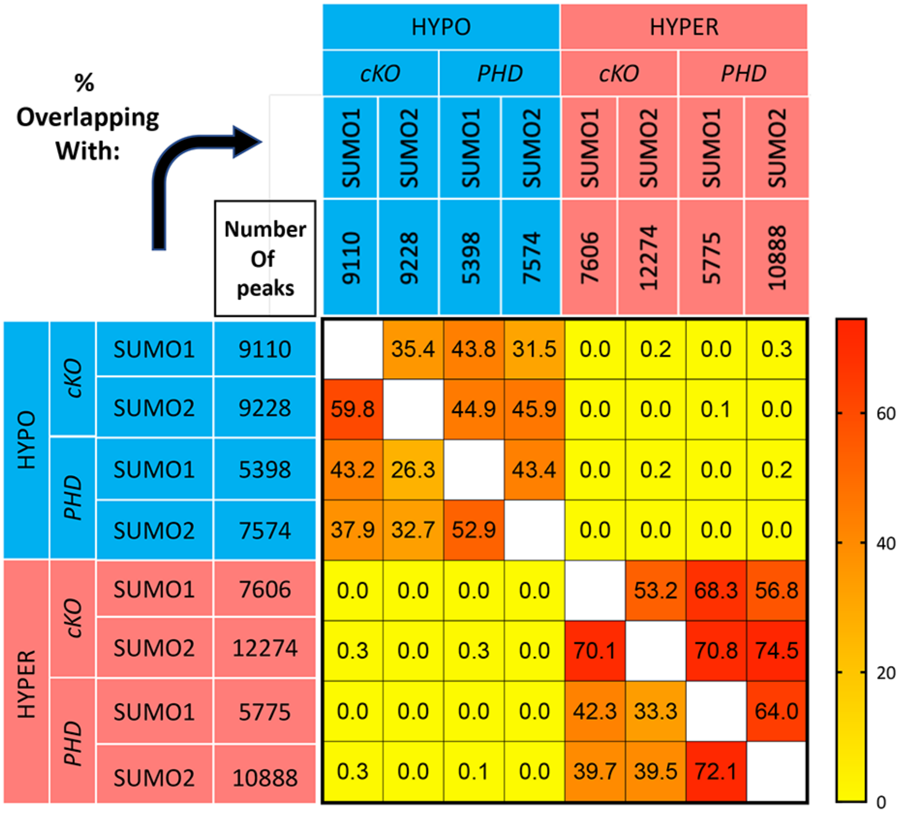


**Fig S20**. Percentage of hyper-SUMOylated ChIP-seq peaks (SUMO1 and SUMO2) in mutant ovaries (*Trim28*$^{cKO}$, *cKO,* and *Trim28*$^{Phd/cKO}$, *PHD*) that overlap with DMRT1and SOX9 peaks from sexual fate reprograming by the testicular transcription factor DMRT1 (*CAG-Stop-Dmrt1-Gfp; Nr5a1-Cre*) in adult ovary [60]. The values in brackets correspond to the number of peaks.

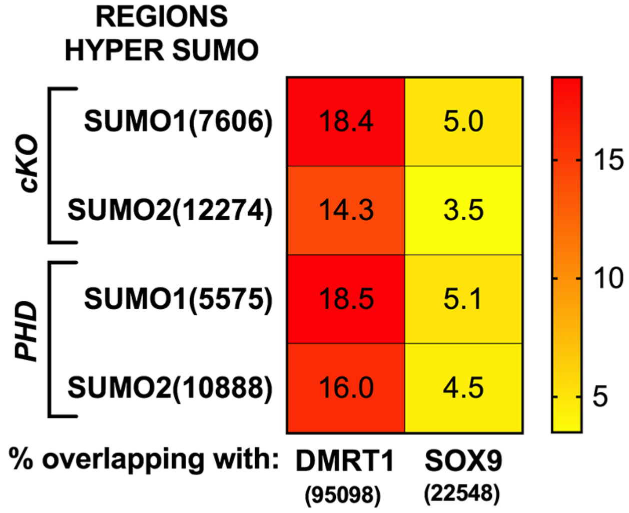

**Fig. S21. a,** Venn diagrams comparing in *Trim28*$^{cKO}$ and *Trim28*$^{Phd/cKO}$ ovaries (*cKO* and *PHD,* respectively) hypo-SUMOylated (left diagram) and hyper-SUMOylated (right diagram) genes.
**b,** Venn diagrams comparing hypo- and/or hyper-SUMOylated genes in *Trim28*$^{cKO}$ (*cKO*, left) and *Trim28*$^{Phd/cKO}$ (*PHD*; right) ovaries. In both mutants, some genes display both hypo- and hyper-SUMOylated peaks (2119 in cKO and 1319 in PHD).

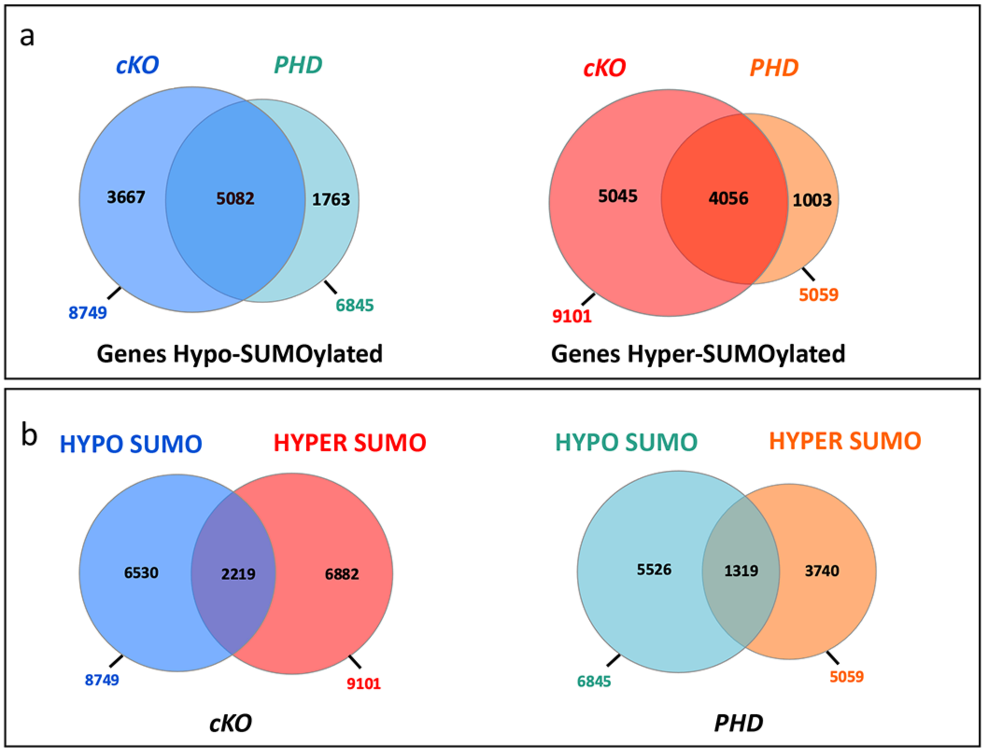

**Fig. S22.** SUMOylation status (SUMO1 and 2) in *Trim28*$^{cKO}$ and *Trim28*$^{Phd/cKO}$ ovaries (*cKO* and *PHD*, respectively) of genes that are downregulated in *Trim28*$^{cKO}$ ovaries. Blue and red backgrounds highlight hypo-SUMOylated and hyper-SUMOylated regions, respectively. Blue and red triangles represent the centre of TRIM28 and FOXL2 peaks, respectively (see fig S11).

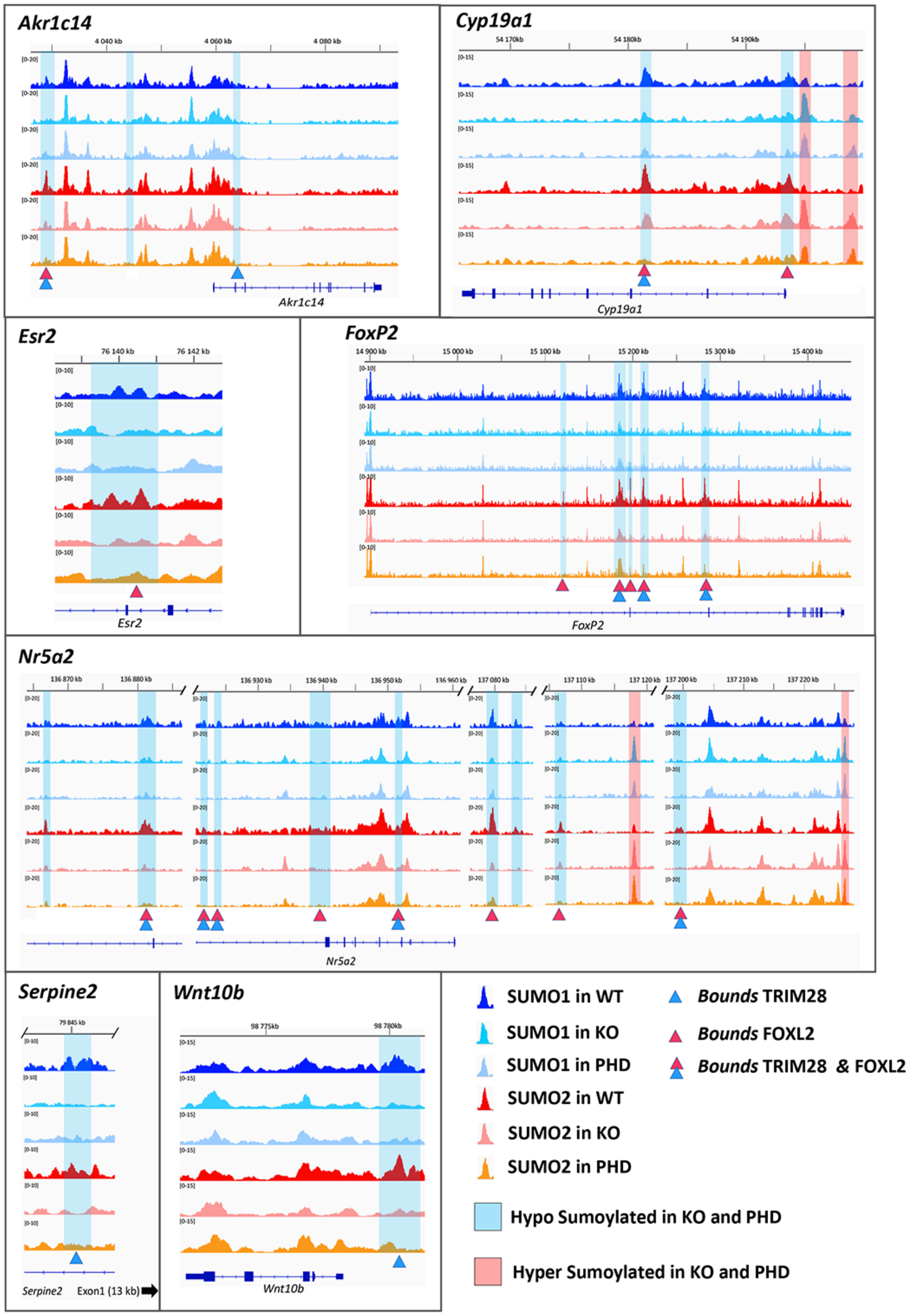

**Fig. S23.** Same as in fig. S22, but for genes that are upregulated in *Trim28*$^{cKO}$ ovaries. Blue and red backgrounds highlight hypo-SUMOylated and hyper-SUMOylated regions, respectively. Blue and red triangles represent the centre of TRIM28 and FOXL2 peaks, respectively (see fig S12).

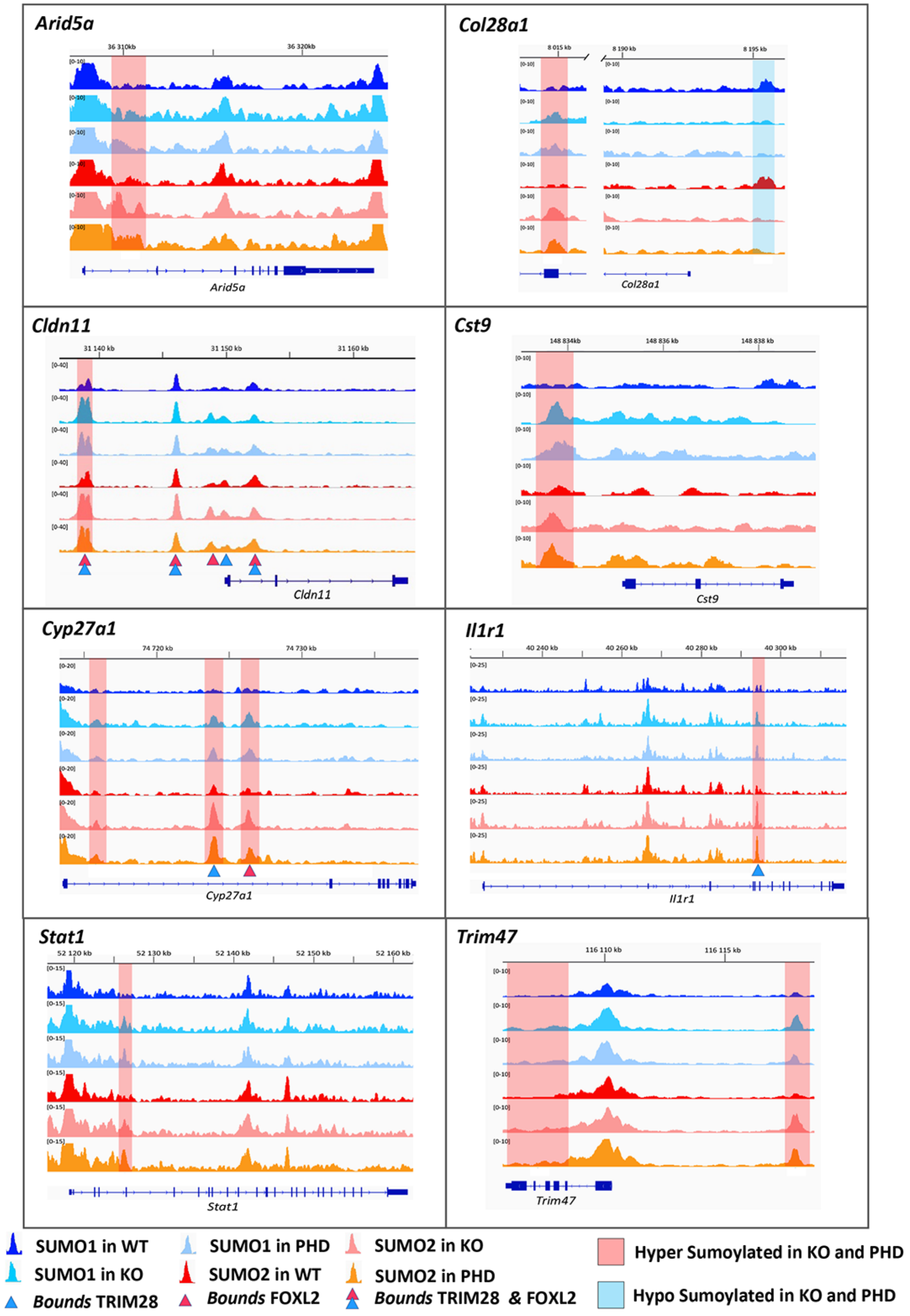
